# Type IV pili-associated secretion of a biofilm matrix protein from *Clostridium perfringens* that forms intermolecular isopeptide bonds

**DOI:** 10.1101/2024.11.04.621531

**Authors:** Sarah E. Kivimaki, Samantha Dempsey, Collette Camper, Julia M. Tani, Ian K. Hicklin, Crysten E. Blaby-Haas, Anne M. Brown, Stephen B. Melville

## Abstract

*Clostridium perfringens* is a Gram-positive, anaerobic, spore-forming, bacterial pathogen of humans and animals. *C. perfringens* also produces type IV pili (T4P) and has two complete sets of T4P-associated genes, one of which has been shown to produce surface pili needed for cell adherence. One hypothesis about the second set of T4P genes is that they comprise a system analogous to the type II secretion systems (TTSS) found in Gram-negative bacteria, which is used to export folded proteins from the periplasm through the outer membrane to the extracellular environment. Gram-positive bacteria have a similar secretion barrier in the thick peptidoglycan (PG) layer, which blocks secretion of folded proteins >25 kD. To determine if the T4P-associated genes comprise a Gram-positive TTSS, the secretome of mutants lacking type IV pilins were examined and a single protein, a von Willebrand A domain containing protein, BsaC (CPE0517), was identified as being dependent on pilin PilA3 for secretion. The *bsaC* gene is in an operon with genes encoding a SipW signal peptidase and two putative biofilm matrix proteins BsaA and BsaB, both of which have remote homology to *Bacillus subtilis* biofilm protein TasA. Since BsaA forms long oligomers that are secreted, we analyzed BsaA monomer interactions with *de novo* modeling. These models projected that the monomers formed isopeptide bonds as part of a donor strand exchange process, in which an N-terminal disordered loop of one monomer intercalates into a beta sheet structure of an adjacent monomer and reforms into a beta sheet with subsequent isopeptide bond formation. Mutations in residues predicted to form the isopeptide bonds led to loss of oligomerization, supporting an exchange and lock mechanism. Phylogenetic analysis showed the BsaA family of proteins are widespread among bacteria and archaea but only a subset is predicted to form isopeptide bonds.

**Importance:** For bacteria to secrete folded proteins to the environment, they have to overcome the physical barriers of an outer membrane in Gram-negative bacteria and the thick peptidoglycan layer in Gram-positive bacteria. One mechanism to do this is the use of a Type II secretion system in Gram-negative bacteria, which has a structure similar to type IV pili and is modeled to act as a piston that pumps folded proteins through the outer membrane to the environment. *Clostridium perfringens*, like all or most all of the clostridia, has type IV pili and, in fact, has two sets of pilus-associated genes. Here we present evidence that *C. perfringens* uses one set of pilus genes to secrete a biofilm associated protein and may be responsible for secreting the main biofilm protein, BsaA. We show that BsaA monomers are, unlike most other biofilm matrix proteins, linked by intermolecular isopeptide bonds, enhancing the physical strength of BsaA fibers.

## Introduction

Bacterial protein secretion to the exterior environment involves transporting the protein across several physical barriers. For Gram-negative bacteria these include the cytoplasmic membrane (CM), a relatively thin peptidoglycan (PG) cell wall and the outer membrane. Some proteins, after secretion through the CM, need to be folded in the periplasmic space, which can prevent them from transiting the outer membrane. To get around this problem, Gram-negative bacteria have evolved a Type II secretion systems (TTSS), which have many proteins similar to those found in Type IV pili (T4P) (Fig. 1 and (1, 2)). TTSS are modeled to function as pistons that use a short pseudopilus to pump folded periplasmic proteins through a protein channel (secretin) in the outer membrane (Fig. 1 and (3, 4)). Protein secretion in Gram-positive bacteria involves transport across the CM and a usually thick PG layer. There are many protein secretion mechanisms for transiting the CM of Gram-positive bacteria, including the Sec system, Twin arginine translocation (Tat), Type IV secretion systems, Esat-6 or type VII secretion system, flagella export apparatus (FEA), fimbrilin (pilus) protein exporter (FPE), and holin-dependent translocation. The Type IV, Type VII and FEA can translocate proteins across the PG layer (5, 6). However, proteins secreted through the CM via the Sec system, used for the majority of proteins, need to fold and, once they do, the PG layer of Gram-positive bacteria acts as a barrier to secretion to the environment because the mesh size in the PG layer is too small to allow diffusion of globular proteins >25 kD, as determined by measurements of the PG from the Gram-positive bacterium *Bacillus subtilis* (7).

**Figure 1.**
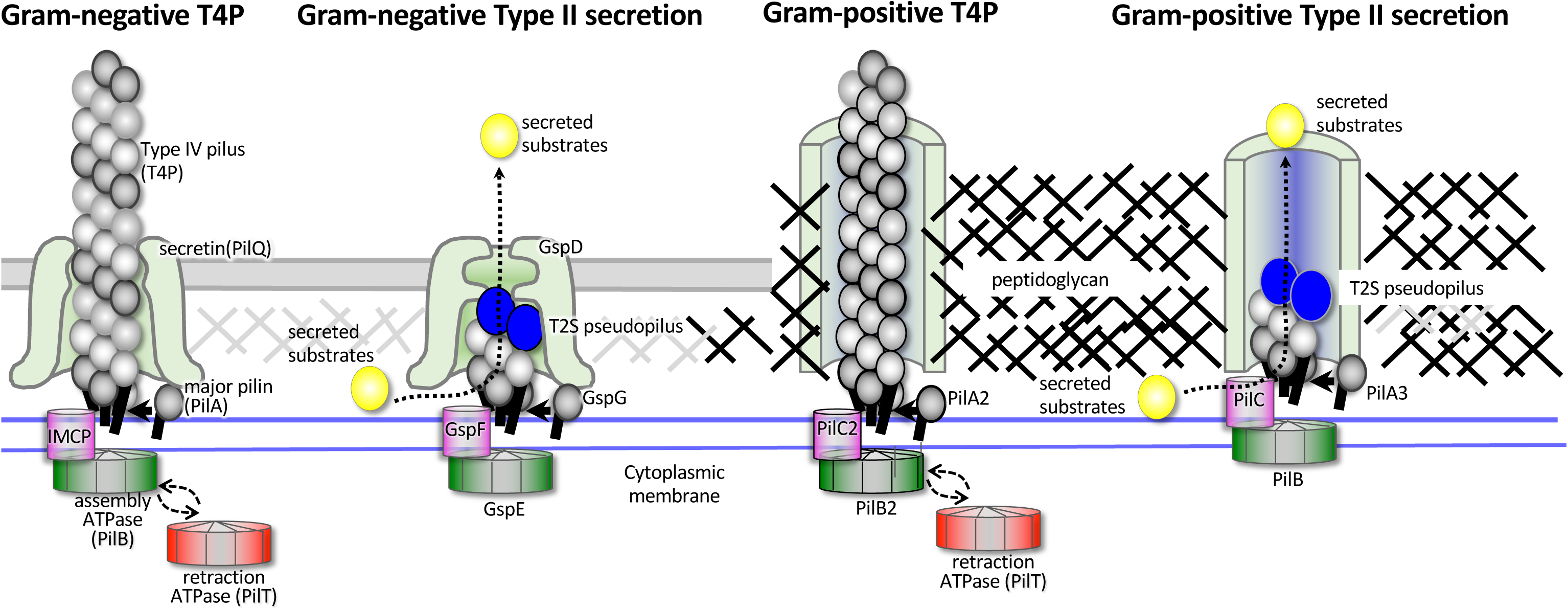
Models showing proposed structures of Gram-positive T4P and Type II secretion systems in comparison to those found in Gram-negative bacteria. Figure modified from Fig. 1 in ref. (9).

One possible mechanism for secretion of folded proteins from the space between the CM and the PG layer in Gram-positive bacteria would be a system analogous to TTSS in Gram-negative bacteria (Fig. 1). Our discovery that T4P systems are ubiquitous and perhaps universal in all the Clostridia (8) provided a means for a potential TTSS to exist in the Clostridia. We hypothesized in a review (9) that in Clostridial species with two separate T4P systems, one of them could function as a TTSS to transport proteins through the PG layer. Multiple species of *Clostridium* do carry two copies of complete T4P systems, including *C. perfringens* (Fig. 2) and *Clostridioides difficile* (9). The core proteins needed for functional T4P usually include a pilin, a pre-pilin peptidase (PilD), an assembly ATPase (PilB), an inner membrane core protein (IMCP, PilC), and inner membrane accessory proteins (IMAP) PilM, PilN, and PilO (9). *C. perfringens* carries multiple pilin-encoding genes, two homologs of PilB, two homologs of PilC and single copies of PilM, PilN, and PilO (Fig. 2), making it plausible that one of these systems could function as a TTSS.

**Figure 2.**
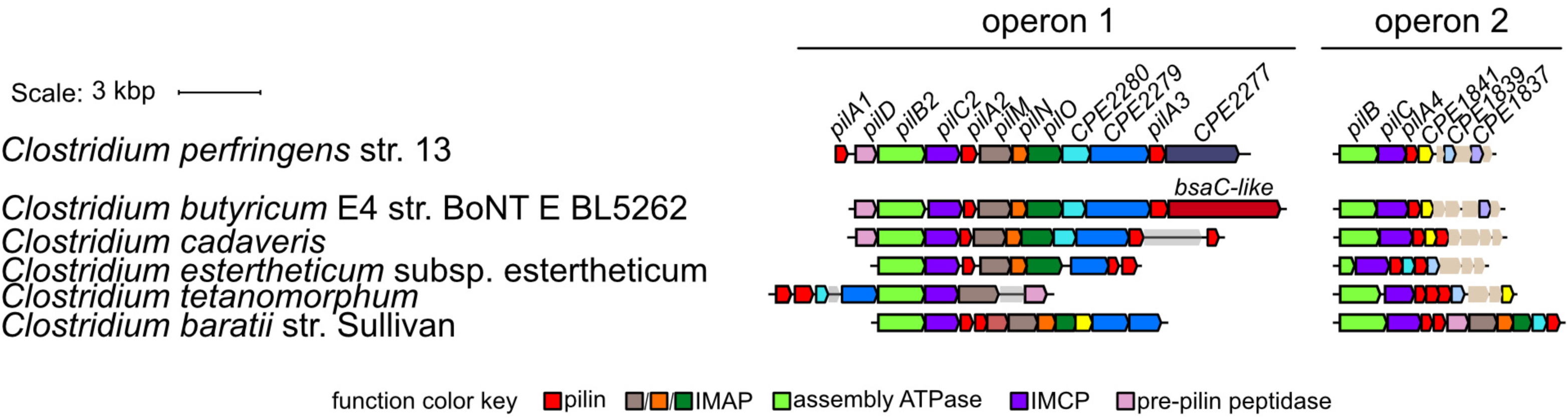
Diagram showing the type IV pili-associated genes in *C. perfringens* strain 13 and other species with two T4P systems. IMCP, inner membrane core protein; IMAP, inner membrane accessory protein.

In 2017 a paper by Obana et al (10) reported that among many other gene transcripts, RNase Y regulated the transcript levels of an operon, *CPE0514-CPE0517*, containing genes encoding extracellular proteins. A later report by Obana et al (11) was published in 2020 in which they examined the regulation and several other features of the *CPE0514-CPE0517* operon that encoded a SipW signal peptidase, two biofilm matrix proteins and a protein with a von Willebrand A domain, the genes of which they designated as *sipW-bsaA-bsaB-bsaC*. BsaA, BsaB, and BsaC are all predicted to have secretion signal sequences (Signal-P 6.0), which suggests they are secreted via the Sec system and the signal sequence is then removed by the SipW signal peptidase. BsaA and BsaB have low levels of sequence homology to the well-studied biofilm matrix protein TasA of *B. subtilis*, (12–15), indicating BsaA and BsaB may also be biofilm matrix proteins. While much of the report of Obana et al. from 2020 concerned the regulation of matrix protein synthesis under different experimental conditions (11), they determined that the BsaA protein was the main component of the *C. perfringens* biofilm matrix. They also discovered some important physical features of BsaA: (1) it required SipW for signal processing and secretion; (2) expression in *E. coli* of a form of BsaA lacking a signal sequence led to spontaneous oligomerization of BsaA, which was resistant to denaturation by SDS or formic acid, indicting the oligomers were held together by very strong intermolecular interactions; (3) a mutant lacking BsaB still formed biofilm matrix oligomers, suggesting it was not important in BsaA oligomerization (11)

These two seemingly disparate areas of research described above overlapped when we undertook experiments to determine if T4P-associated pilin proteins were involved in protein secretion. Prior to the published observations of Obana et al, we discovered that the von Willebrand A domain protein, CPE0517, designated as BsaC by Obana et al. (11), required the PilA3 pilin for secretion (see Results section, below). This demonstrated for the first time that a Gram-positive bacterium required T4P proteins for secretion, providing strong support for our hypothesis that a TTSS-like system is functioning in a Gram-positive bacterium. This finding led us to test whether the main biofilm matrix protein, BsaA, also required T4P-associated proteins for secretion. While our findings indicate that several T4P proteins are needed for secretion of BsaA oligomers, these experiments also led to our discovery that the oligomers of BsaA observed by Obana et al and our group involved formation of isopeptide bonds between BsaA monomers. We then characterized the nature of these bonds and conservation of isopeptide bond formation based on phylogeny of the BsaA/TasA family of proteins. Identifying a potential TTSS-like system in a Gram-positive bacterium and the novel finding of isopeptide bond formation in a well-studied family of biofilm matrix proteins will lead to significant advances in both of these emerging fields of microbial cell biology.

## Results

### Secretome analysis of a *pilA3* mutant indicates a 70 kD protein with a Von Willebrand domain A (CPE0517, BsaC) is absent

To test our hypothesis that T4P proteins were needed for protein secretion, we examined the secretome of strains with in-frame deletions in genes encoding Type 4 pilins We screened the secretome of in-frame deletion mutants in four pilin-encoding genes, *pilA1*, *pilA2*, *pilA3*, *pilA4* for missing proteins, indicating they may be dependent on the pilin for secretion. Only the *pilA3* mutant showed evidence of a distinct missing band in the secretome and this band was restored by complementation of a WT copy of the *pilA3* gene (Fig. 3A). The band was excised from a gel and subjected to trypsin digestion and analysis by mass spectrometry. Peptide fragments matched those of the CPE0517 (BsaC) protein from the *C. perfringens* strain 13 genome. The protein has an unknown function but does contain a von Willebrand A domain, which can potentially function in protein-protein interactions with host cells. Western blots using anti-BsaC antibodies showed a similar loss of CPE0517 secretion by the *pilA3* mutant and complementation by expression of a WT copy of the *pilA3* gene (Fig. 3B). This gene is the fourth gene in a four gene operon containing a type I signal peptidase (CPE0514, SipW) and two genes (*CPE0515-CPE0516*, *bsaA-bsaB*) encoding proteins with similarity to the TasA protein in *B. subtilis* (Fig. 3C).

**Figure 3.**
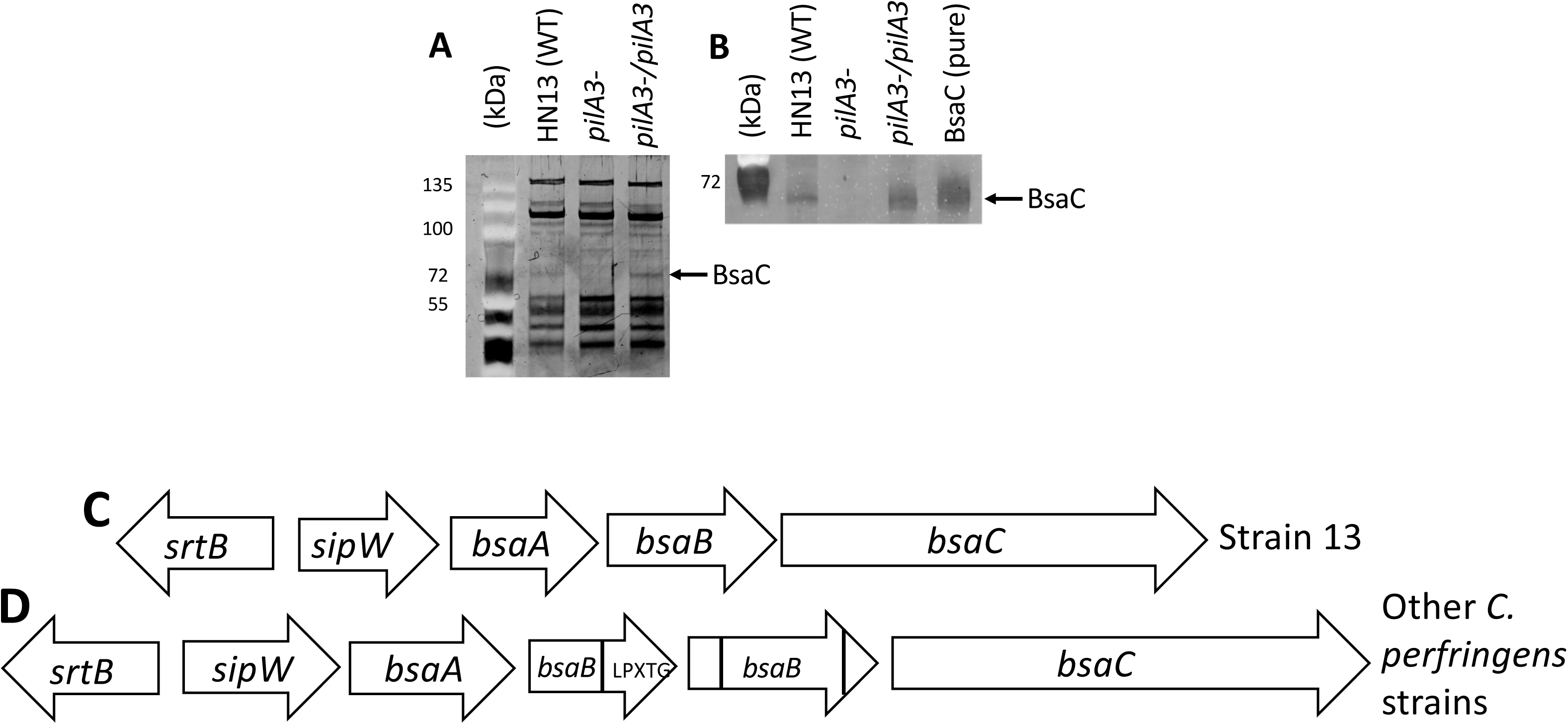
(A). SDS-PAGE gel showing the proteins in the secretome of the respective strains. The gel was stained with the fluorescent protein dye Sypro-Ruby, the colors inverted to make the protein bands in the samples appear dark but the pre-stained molecular weight markers appear as bright bands. **(B).** Western blot showing the levels of secretion of the BsaC protein in different strains. The secondary antibody was fluorescently tagged for visualization. Purified BsaC was added in lane 5 as a positive control for detecting BsaC. **(C).** The gene synteny of the *bsa* operon and adjacent sortase-encoding gene in *C. perfringens* strain 13. **(D).** The gene synteny of the *bsa* operon and adjacent sortase-encoding gene in *C. perfringens* strain ATCC 13124 and most other strains of *C. perfringens* that were observed. The sections of the *bsaB* gene from strain 13 that were homologous to those in strain ATCC 13124 are labeled, along with the location of an LPXTG sortase dependent sequence in one of the *bsaB*-encoding genes.

To determine if other Type IV pilus (T4P) related genes were needed for secretion of BsaC, we expressed the *bsa* operon in a bank of in-frame deletion mutants encompassing the gene known be required for synthesis and assembly of T4P (Fig. 2) using a vector we developed containing a lactose-inducible promoter (16). The amounts of secreted BsaC were measured using quantitative western blotting of TCA precipitated secreted proteins. We noted variation in day-to-day results with the WT strain and, since these were western blots on TCA precipitated protein from culture supernatants, we normalized all of the data from mutant strains to that of the WT strain on the same blot (see legend in Fig. 4). Using quantitative densitometry on western blots of the secreted oligomers, we measured statistically significant decreased secretion of BsaC in the following mutants: *pilA3*, *pilB2*, *pilC1*, *pilD*, *pilM*, *pilN* and *pilO* (Fig. 3). The proteins encoded by these genes have the ability to form a complete Type IV pilus. Mutations in the *pilT* gene (encodes a T4P retraction ATPase), *cpe2277* (encodes a protein of unknown function) and *cpe2279* and *cpe2280* (encode minor pilins) did not affect the levels of secretion of BsaC.

**Figure 4.**
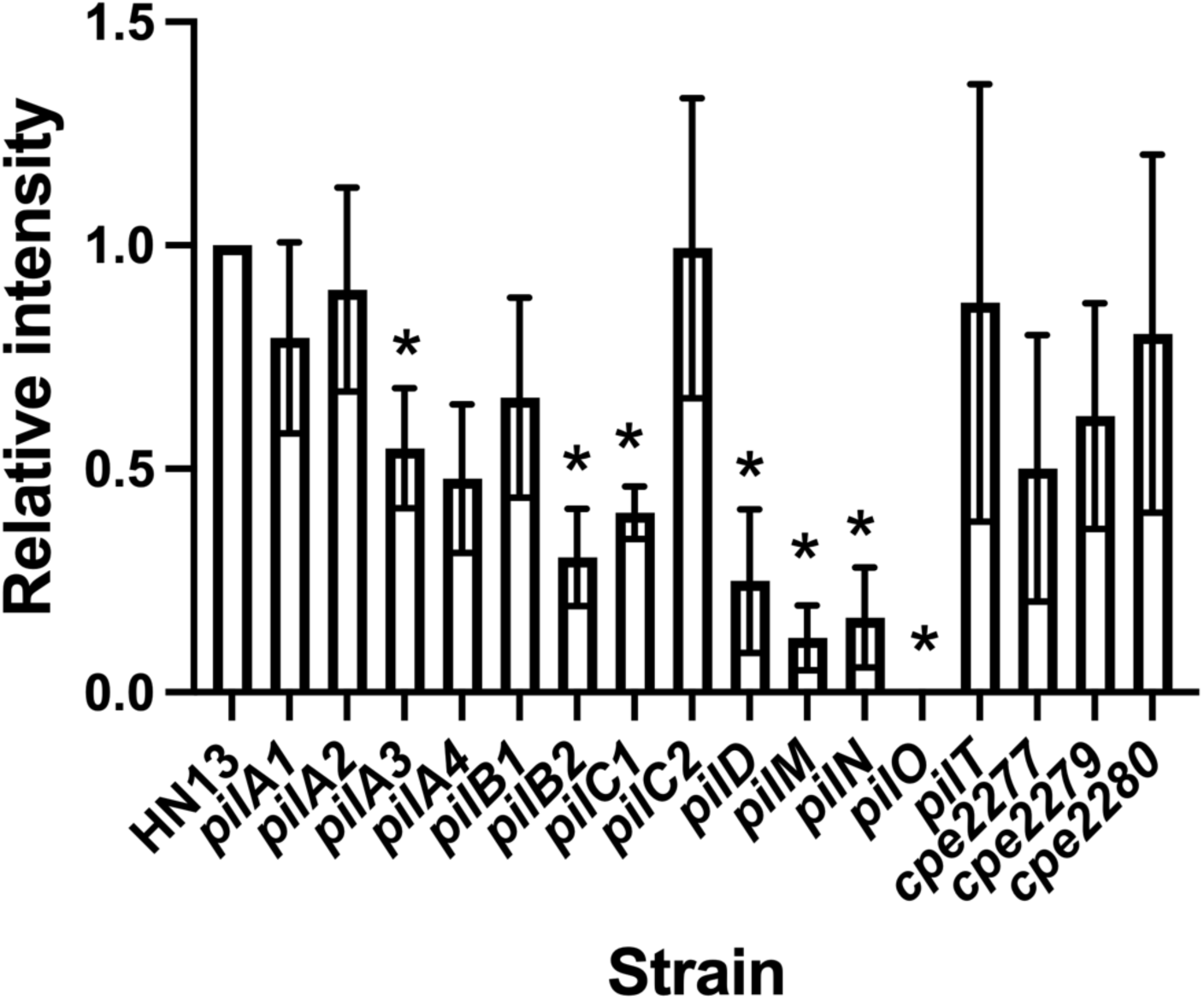
Efficient secretion of the BsaC protein requires T4P proteins. Results from densitometry analysis of western blots from the secretome of strains with in-frame deletions of T4P-encoding genes. Each strain contained plasmid pHLL65 for regulated expression of the *bsa* operon (*sipW-bsaA-bsaB-bsaC*) in the wild type (HN13) and mutants deleted for T4P-associated genes. After expression, secreted proteins were concentrated from culture supernatants using TCA and quantitative western blotting was performed. The role of each gene product in pilus function is shown in Fig. 2. For each experiment, the relative fluorescence intensity of the wild-type strain (HN13) was set at 1.0 and the relative fluorescence intensity for each other strain in the same experiment was then set relative to that value. Each strain was tested at least three times with triplicate samples for each experiment and the mean and SEM are shown. All the experimental runs were combined, and one sample t and Wilcoxon tests were run to determine if the mutant strains were statistically different from 1. Asterisks indicate strains that were different (*P* < 0.05) from strain HN13 (i.e., 1).

A major difference in the structures of T4P and TTSS is the type IV pilus extends beyond the outer membrane (Gram-negative) or PG layer (Gram-positive, (8)) but in the TTSS in Gram negative bacteria the pilus does not (Fig. 1 and (2)). Our original model for a TTSS in Gram-positive bacteria is that the pilus (i.e., PilA3) does not extend past the PG layer (Fig. 1). To test this model, we exposed *C. perfringens* cells to rabbit anti-PilA3 antibodies and a fluorescently tagged goat anti-rabbit secondary antibody and found that we could detect binding localized to the poles of the cells (Fig. S1). This refined the location of the PilA3 pili to the poles, and that they extend far enough out past the PG layer to be exposed to antibodies, which suggests the model shown in the right most panel in Fig. 1 may need to be modified to show extension of the pilus to the outside environment.

To determine if there are generalized secretion defects in T4P mutants, we examined whether each mutant in the T4P-associated genes was able to secrete another protein, the phospholipase-sphingomyelinase, PLC. Secretion of PLC can be easily detected by a ring of precipitation around a colony using egg yolk agar, and we found that none of the T4P mutants showed a detectable decrease in PLC secretion (Fig. S2), suggesting the secretion defects seen with BsaC were not due to generalized secretion defects.

### Expression of the *bsa* operon leads to synthesis of extracellular biofilm matrix material

Since the *bsa* operon encodes proteins with similar synteny and predicted structural homology to the *sipW-tasA-tapA* biofilm matrix operon in *B. subtilis* (15), we placed the entire operon (*sipW-bsaA-bsaB-bsaC*) under control of the lactose inducible promoter for high level expression. The cells produced large quantities of an extracellular gel like material (Fig. S3A), which, after staining with crystal violet, appeared as amorphous material between the bacterial cells (Fig. S3B). A His_6_ tag was added to the C-terminus of the BsaA protein and, when the operon was expressed, a distinct high molecular weight ladder of BsaA was visible in anti-His_6_ antibody western blots after running on SDS-PAGE gels using samples boiled in SDS-PAGE buffer (Fig. S3C). This suggested BsaA was in very heat-stable oligomers. This was in agreement with the results of Obana et al who found that BsaA was the major component of the biofilm matrix and that BsaA oligomers were stable after boiling in SDS-PAGE buffer and resistant to dissociation by high levels of SDS and formic acid (11).

### Efficient secretion of the BsaA oligomer is dependent on some T4P-associated genes

Since it was possible that, like BsaC, BsaA in its oligomer form was secreted with the aid of T4P-associated proteins, we expressed a clone containing just the *sipW* and *bsaA* genes in which a FLAG-tag was added to the C-terminus of BsaA for quantification in western blots using densitometry. Only *sipW* and *bsaA* were included to simplify the analysis of the resulting western blots. An example of a typical blot is shown in Fig. 5A. Using this method, we found that the *pilA1*, *pilB1*, *pilB1/pilB2*, *pilN* and *pilO* mutants exhibited decreased levels of BsaA oligomers in the supernatants (Fig. 5B). However, since the genes required for secretion lacked an inner membrane core protein (*pilC*) component and pre-pilin peptidase, which are essential for T4P functions (17), the results were not definitive.

**Figure 5.**
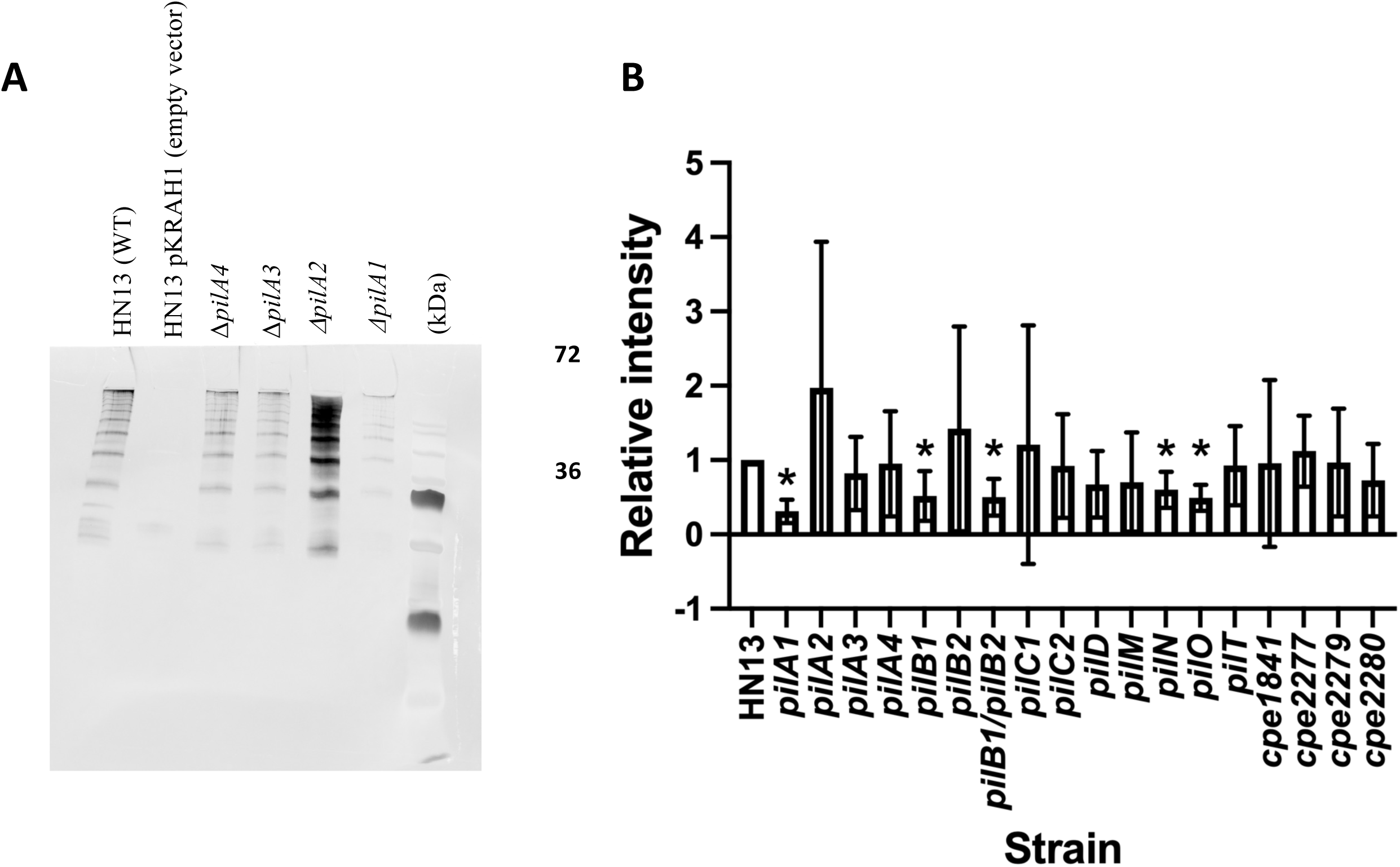
Use of western blots of secretomes to determine if T4P genes play a role in BsaA oligomer secretion. After culture in BHI medium of strain HN13 with pSK4 (*sipW-CPE0515*-FLAG), proteins were precipitated from culture supernatants by using TCA. **(A)** Sample blot showing the presence of oligomers in the wild-type and some mutant strains. Densitometry was used to determine the intensity of the fluorescent signal for all the oligomer forms that were present. **(B)** Summary of results from all the T4P-associated mutants. Each strain was tested at least six times in independent experiments and the mean and SEM are shown. For each experiment, the relative fluorescent intensity of the wild-type strain (HN13) was set at 1.0 and the relative fluorescence intensity for each other strain in the same experiment was then set relative to that value. All the experimental runs were combined, and one sample t and Wilcoxon tests were run to determine if the mutant strains were statistically different from 1. Asterisks indicate strains that were different (*P* < 0.05) from 1.0 (i.e., strain HN13).

### Models of BsaA oligomers predicts an isopeptide bond is formed linking adjacent monomers

BsaA forms very heat and SDS resistant oligomers ((11) and Figs. S3C and 5A). To investigate the structural nature of the interactions between monomers of BsaA that leads to this stability, a structure would need to be developed of a homomultimer. Due to the difficulty of solving the structure of long, thin chains by experimental methods, computational modelling was used. Models were initially created using Phyre2 (18), with the top model being an x-ray crystal structure of TasA from *B. subtilis* (13). That structure, and homology models from that structure, were monomeric, and not useful for determining intermolecular interactions between monomers. AlphaFold Multimer was then used to create a multimeric model to determine inter-chain interactions. Using ColabFold 1.3.0 (19), which is based on AlphaFold 2.2.0 (20), a homotrimeric structure of BsaA was created (Fig. 6A). The SipW-specific signal sequences (residues 1-27) were included in the initial model, but only formed disordered loops which do not have any notable interactions. Removing the signal sequences did not change the overall structure of the monomers so this was done for subsequent analyses.

**Figure 6.**
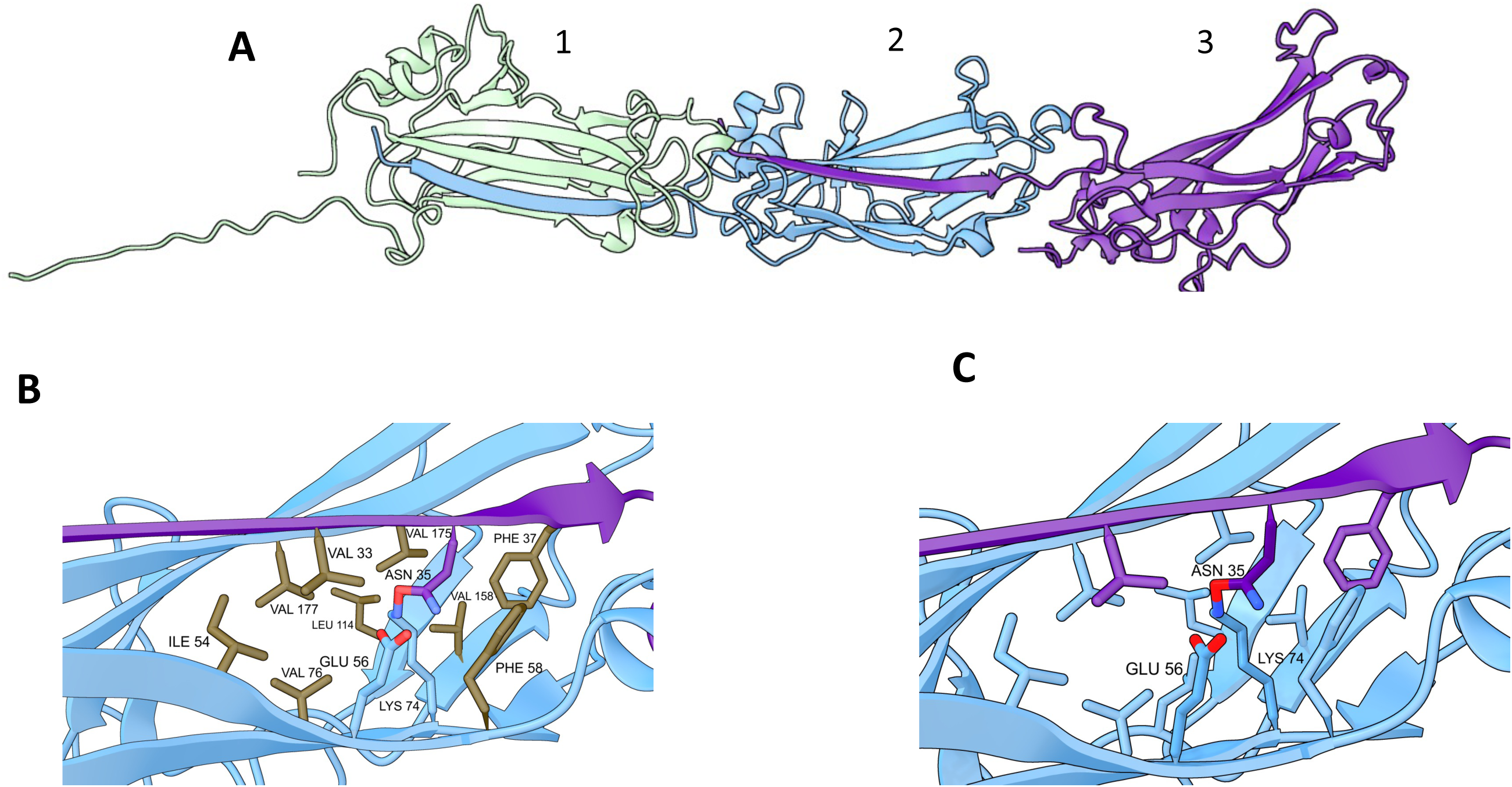
(A) AlphaFold model of a homotrimer of BsaA in cartoon representation. Each independent monomer is colored separately. Note that residues 28-38 of a monomer incorporate themselves into the beta strands of an adjacent monomer. **(B)** Close-up view of interdigitated beta strands from two monomers in cartoon representation, showing relevant side chains as sticks colored by hydrophobicity. N35 and the giving chain are colored purple, E56, K74, and the receiving monomer are colored cyan. Side chains forming the hydrophobic pocket are colored brown. The bond created by AlphaFold between N35 and K74 is between an oxygen and nitrogen due to limitations in AlphaFold; in reality, an amide bond would form between the side chains with ammonia as the leaving group. **(C)** Close-up view of the N35-E56-K74 catalytic triad, showing the close proximity of the catalytic E56 residue to the bond forming N35 and K74. All images were created with ChimeraX v1.2. (43, 44).

In monomeric structures, residues 28-38 would extend out as disordered loops (Fig. 6A, monomer 1). In the multimeric structures, these residues incorporated themselves into the beta sheet structure of adjacent monomers (Fig. 6A). Being fully incorporated into a nearby structure indicates that residues 28-38 provide specific interactions with the receiving monomer, providing the basis of a mechanism for which oligomers of BsaA can be created. For example, V33 and F37 of the interdigitated strand become part of the hydrophobic core of the receiving monomer (Fig. 6B). The interactions between specific side chains of residues from the interdigitated strand and the receiving monomer were investigated further for less standard interactions.

The *de novo* created structures indicate a bond between two side chains: N35 of the interdigitated strand and K74 of the receiving monomer (Fig. 6C). At first, this was thought to be an error in AlphaFold’s model generation, since the chemistry would not be possible. The bond was created between the oxygen of N35 and the nitrogen of K74, resulting in three bonds in the oxygen. However, this bond was created in nearly every model generated, so it was investigated further. A crosslink between asparagine and lysine is a type of isopeptide bond, which has been characterized through a few different mechanisms and can be found as an intermolecular crosslink, as in ubiquitylation (21), or as an intramolecular crosslink found in Gram-positive pili (22). In Gram-positive pili, this bond is formed between an asparagine and a lysine, autocatalyzed by a glutamate within the hydrophobic core of the protein. Residues equivalent to these are found surrounding the bond in BsaA (Fig. 6C). The hydrophobic region surrounding the N35-K74 pair (Fig. 6B, brown colored residues) is believed to be necessary to help the catalytic cross-linking reaction (23).

It is unclear why this bond was created by AlphaFold. In the documentation of AlphaFold Multimer (24), there are no mentions of predicting crosslinks. One potential reason is that it generated the side chains so close to each other there was a steric clash, but the rest of the model was so favorable the structure was created regardless. Models created in ColabFold have an option to perform energy minimization using Amber. For these models, this was not performed. Recreating this model while enabling energy minimization eliminates the bond every time. This is expected given the incorrect chemistry of the bond. AlphaFold3 (25) does not include a minimization step, and, like AlphaFold2, creates the isopeptide bond.

This amyloid-like fold and interdigitation is seen in other Gram-positive bacteria, such as TasA in *B. subtilis*. A cryo-EM structure has been created for a chain of TasA (12), which demonstrated how hydrophobic interactions between donor and acceptor monomers drive interdigitation. No crosslinks were found or mentioned, and the required triad of N-E-K residues is not present in the structure (PDB 8aur). Therefore, the isopeptide bonds in BsaA oligomers would be the first example of an intermolecular bond of this mechanism found in this family of proteins.

### Mutations in isopeptide bond forming residues abolish oligomerization of BsaA

The AlphaFold Multimer models predicted an isopeptide bond was formed between N35 and K74 from adjacent residues in a BsaA oligomer. To test this computational prediction, we made point mutant substitutions of each residue to alanine, forming N35A and K74A mutants, along with a double N35A/K74A mutant. His-tagged versions of these mutant forms were expressed along with *sipW* and the cell extracts and culture supernatants were examined for the presence of the mutant proteins in oligomeric form. Mutating N35 or K74 to alanine residues resulted in a loss of oligomers in the supernatants and the appearance of monomers of the K74A mutant but not of the N35A mutant, suggesting the N35A version was not effectively released by the cell (Fig. 7A). Monomeric forms of both mutants appeared in the cell extract (Fig. 7A). In some experiments, small amounts of dimers and higher mol wt forms of BsaA appeared in the cell pellet, but the large majority were in monomeric form, which was absent in the WT form of BsaA (Fig. 7A), perhaps due to the stability of oligomers even in the absence of an isopeptide bond. To test this, we suspended cell pellet extracts from BsaA-His_6_ and N35A-His_6_ in SDS-PAGE buffer and heated at 95 C for 0, 10, 20 and 40 min before running the samples on SDS-PAGE gels and performing western blots. The unheated samples were incubated for 40 min at room temp as a control. The N35A-His mutant maintained stable oligomers in SDS-PAGE buffer and after being run through an SDS-PAGE gel in the absence of heating, but heating the samples for 10-40 min led to dissociation of the oligomers, in contrast to the WT BsaA protein, which was not affected by heating (Fig. S4). This suggests that N35, and therefore the isopeptide bond, is essential for forming detergent and heat resistant oligomers, and that the N35 monomer-monomer interactions are stable even in the presence of 2% SDS (the concentration in the SDS-PAGE buffer), which is in contrast to oligomers formed by the *B. subtilis* TasA protein, which were dissociated by SDS (14).

**Figure 7.**
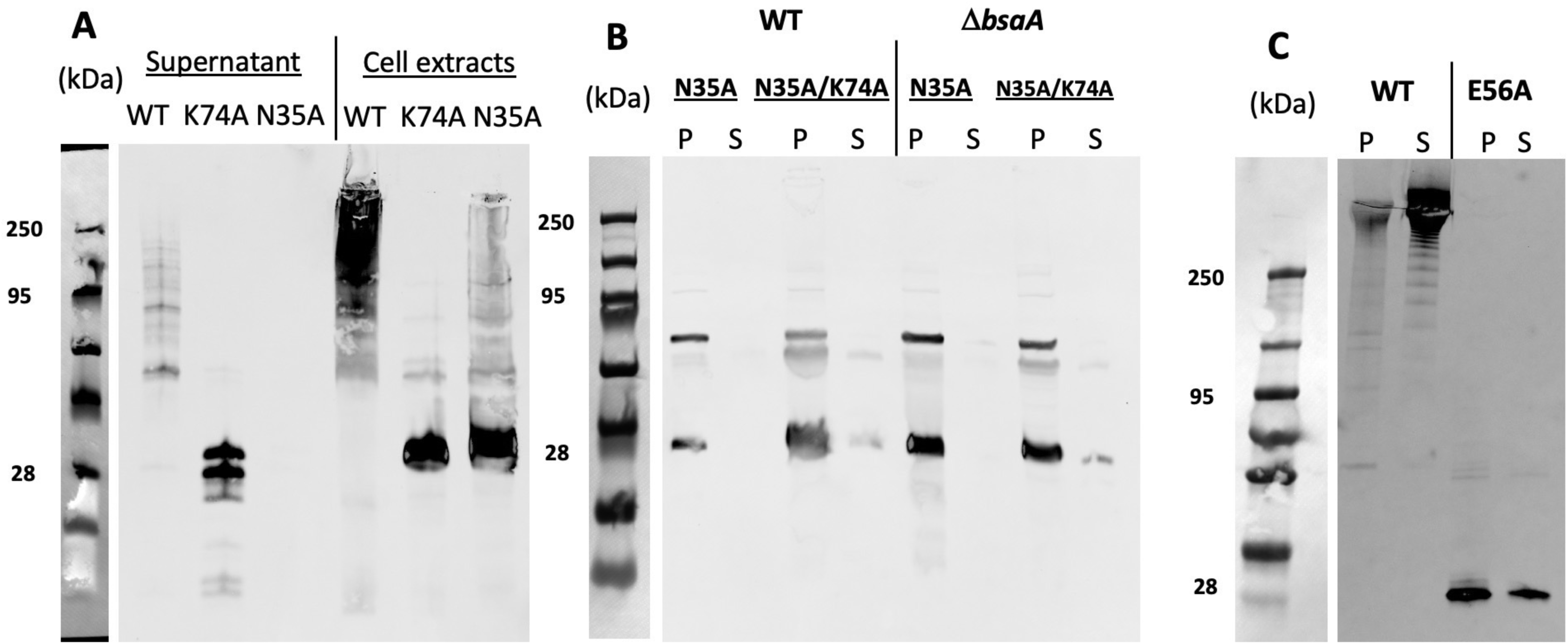
Western blots showing loss of oligomer formation in BsaA with mutations in residues involved in isopeptide bond formation**. (A)** Oligomer formation seen in TCA precipitated culture supernatants or in the disrupted whole cell extracts of strains producing BsaA with single mutations. **(B)** Comparison of BsaA oligomer formation in the N35A and N35A/K74A mutants in a wild-type background (left side) and one in which the chromosomal copy of the *bsaA* gene was deleted. Note that the chromosomal *bsaA* mutation did not change the oligomerization levels of the mutants. **(C)** Western blot showing mutation of the E56 residue to Ala abolished oligomerization of BsaA. P, disrupted cell pellet; S, TCA precipitate of culture supernatant.

We also tested the N35A/K74A mutant and observed that it showed a similar loss of oligomerization as did the N35A mutant (Fig. 7B, left four lanes). These experiments were all done in strain HN13 (WT) background where there was an intact *bsaA* gene. To determine if this potential source of BsaA, even though it was not His-tagged, affected the results we observed with the mutants, we expressed the N35A and N35A/K74A mutants in a strain with an in-frame deletion of the *bsaA* gene, but did not observe any differences to the pattern seen in the strain with an intact *bsaA* gene (Fig. 7B, right four lanes), suggesting the chromosomal source of BsaA was not forming oligomers in combination with the N35A and N35A/K74A mutant forms of BsaA.

### The Glu56 residue of BsaA is essential for efficient isopeptide bond formation

Spontaneous or self-catalyzed isopeptide bond formation is catalyzed by an acidic Glu or Asp residue located within the hydrophobic pocket where the Asn and Lys residues are located (23). In AlphaFold 2 BsaA models of dimer interfaces, Glu56 is predicted to be in the precise location to catalyze bond formation (Fig. 6C). To test if this was actually occurring, we constructed an E56A substitution in the BsaA protein and examined its ability to oligomerize by using western blotting on cell extracts and concentrated supernatants from the mutant and WT strain (Fig. 7C). The E56A mutant showed a clear lack of oligomer formation and an accumulation of the monomeric form of the protein which was not observed with the WT form of BsaA (Fig. 7C), providing strong evidence that Glu56 is responsible for catalyzing isopeptide bond formation in BsaA.

### A FAST fusion to BsaA displays surface exposed localization

To investigate the cellular location of the BsaA oligomers, a FAST gene fusion to the *bsaA* gene was constructed and, along with the *sipW* gene, was placed under control of the lactose-inducible promoter. The FAST protein binds a soluble dye which, upon binding, exhibits a large increase in fluorescence (26–28) allowing localization of the fusion proteins. We expressed the *sipW-bsaA*-FAST fusion genes and exposed the cells to two different dyes, Coral, which is membrane permeable, and Amber-NP which is impermeable to membranes (both obtained from The Twinkle Factory). To test if the Coral and Amber-NP permeability profiles were working in *C. perfringens* membranes, we expressed a *pilT*-FAST gene fusion, which should be located only in the cytoplasm, and exposed them to the Coral and Amber-NP dyes. Consistent with the predicted cytoplasmic location of the *pilT*-FAST fusion, only the Coral dye showed significant fluorescence after treatment (Fig. 8A). However, when the *bsaA*-FAST fusion was expressed along with *sipW*, both dyes were able to bind and fluoresce (Fig. 8B), indicating the BsaA-FAST fusion was competent for secretion in *C. perfringens* and was located on the surface of the bacteria.

**Figure 8.**
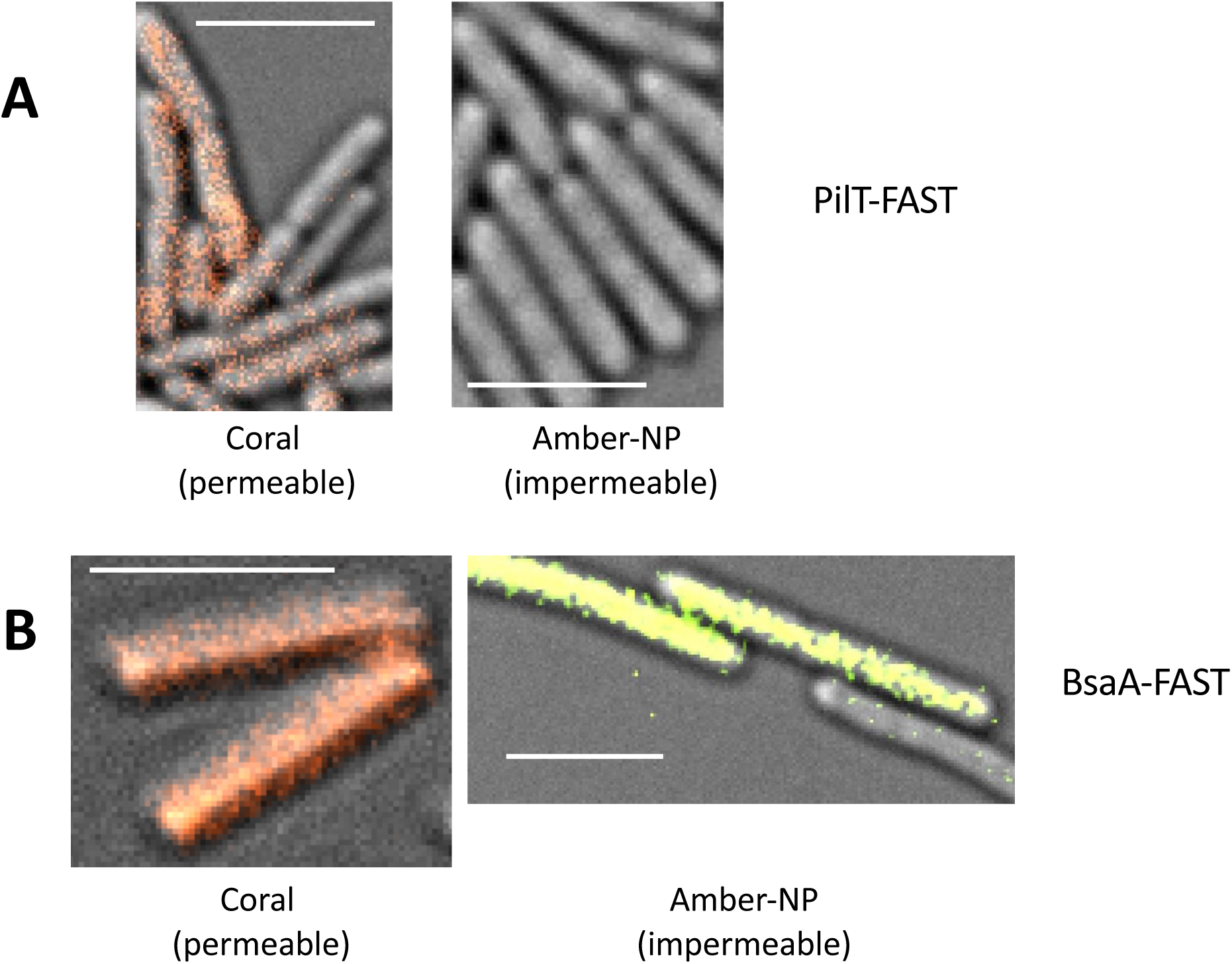
BsaA oligomers are found on the bacterial surface. **(A)** Expression of a FAST protein fusion to the cytoplasmic PilT protein (pSM413) only shows fluorescence with the membrane permeable Coral dye. **(B)** A BsaA-FAST fusion protein (pSM412) fluoresces in the presence of both a membrane permeable dye (Coral) and a membrane impermeable dye (Amber-NP), indicating it is surface exposed to these dyes. Scale bars = 5 μm.

### BsaA oligomers are anchored to the membrane and not the peptidoglycan in strain HN13

Because the BsaA-FAST fusion experiments indicated BsaA oligomers were located on the cell surface and SEM images of the BsaA matrix proteins suggested they were in the form of fibrils associated with the bacterial surface (11), we wanted to determine if the oligomers were embedded in the cytoplasmic membrane or covalently linked to the PG layer. To test this, we expressed *sipW-bsaA*-His_6_ constructs for the WT and each mutant in our lactose-inducible promoter system and then boiled the cells in 10% SDS for 30 min, followed by digestion of the PG matrix by lysozyme. Our experimental approach was based on the hypothesis that if BsaA was anchored in the membrane, boiling in SDS would release it from the cell but if it was covalently anchored to the PG, it would be freed only after digestion with lysozyme. We tested the WT, N35A, K74A and N35/K74 mutants and found that all of the long wild-type oligomers were released after the SDS boiling step, while the cells appeared to retain a small number of monomers and dimers of BsaA (Fig. S5). However, after treatment with lysozyme, the monomers and dimers were no longer detected. For the mutants that can’t oligomerize, the monomer/dimers were retained after boiling in SDS, but not after lysozyme treatment. We interpret these findings as indicating the oligomers are anchored to the membrane but after SDS boiling they are released. A small number of monomers and dimers remain trapped inside the sacculi produced by boiling in SDS but lysozyme treatment releases these from the sacculi. However, they are then too dilute to detect on western blots. This is consistent with an analysis of the *bsaA* and *bsaB* protein sequences, both of which lack a recognizable LPXTG motif, despite the presence of a sortase gene, *srtB*, adjacent to the operon (Fig. 3C).

### BsaB, an ortholog of BsaA, forms dimers but not long oligomers

Previous results by Obana et al (11) showed a *bsaB* mutant still made significant amounts of the BsaA oligomers but they did not test to see if BsaB was incorporated into BsaA oligomers. Therefore, we added an HA-tag to the BsaB protein and expressed the *sipW-bsaA-bsaB*-HA-*bsaC* genes in our inducible system and examined the culture supernatants for BsaB-HA using western blots. Based on the predicted molecular weight of the BsaB protein, we detected BsaB-HA only in a monomeric and a higher molecular weight form which could be either a homodimer or a BsaA-BsaB heterodimer (Fig. S6), but it is not covalently attached to the longer BsaA oligomers.

### Phylogeny of isopeptide bond formation in the TasA family

Since a BlastP search with BsaA only identifies homologs in closely related Clostridial species, an iterative search with profile hidden Markov models was employed. This analysis resulted in the identification of 2,172 proteins, including the previously described TasA from *B. subtilis* and CalY from *Bacillus cereus* (Fig. 9). Of these, 632 hits belong to the “peptidase M73, camelysin” (InterPro: IPR022121) family; so-called because of the likely misidentification of CalY as a peptidase. In addition to the use of probabilistic models, the relatedness of these proteins is evident at the structural level. Like other family members, BsaA is predicted to adopt a β-sheet–rich Ig-fold-like structure and is predicted to participate in donor-strand exchange (Fig. 6), as described for TasA (12). These analyses suggest that while the structures of these proteins and homopolymer assembly are highly conserved, sequences are highly divergent.

**Figure 9.**
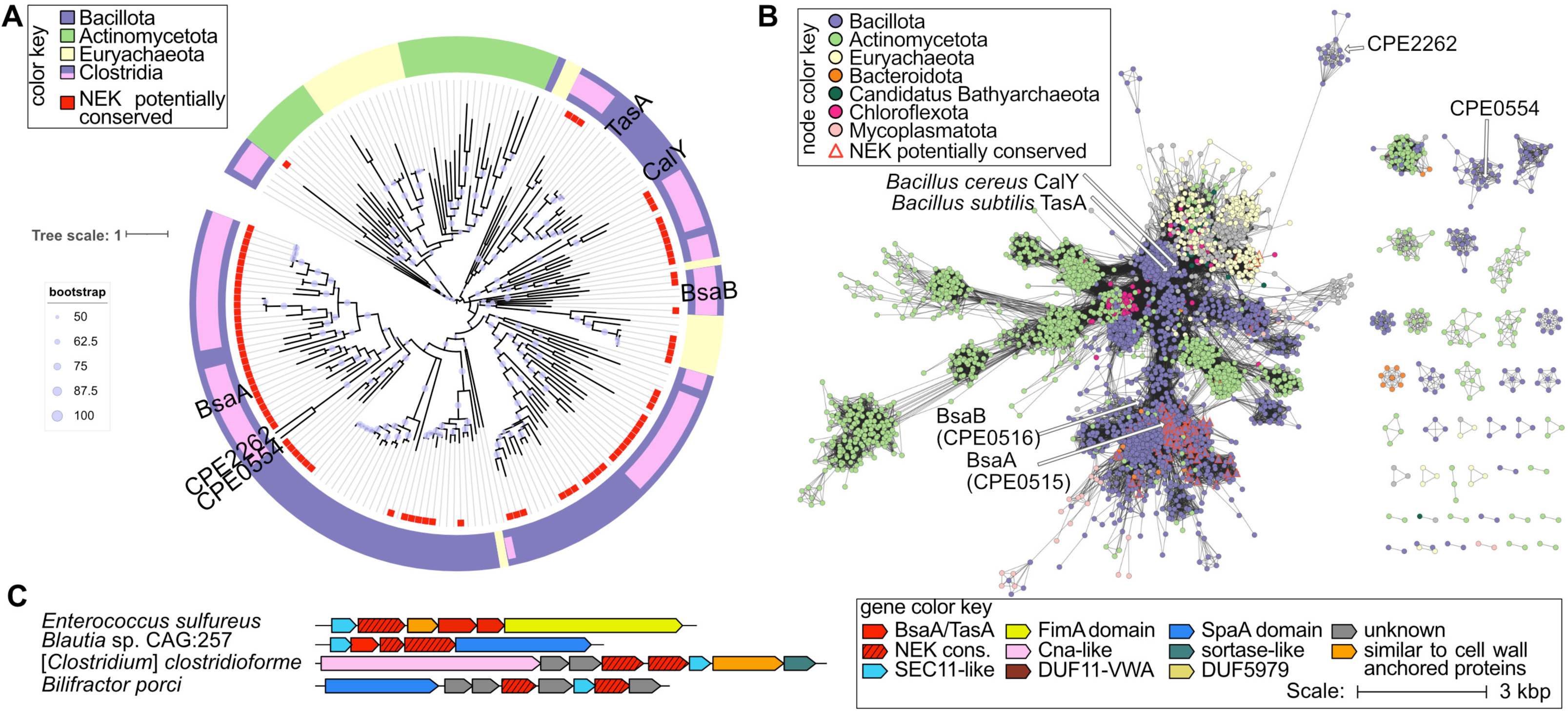
BsaA phylogeny. (**A**) A consensus tree of representative BsaA-like proteins. Leaf labels are colored by taxonomy according to the color key. BsaA, BsaB, and two other BsaA-like proteins identified in *C. perfringens* are labeled, as are TasA from *B. subtilis* and CalY from *B. cereus*. Bootstrap values are represented as scaled circles according to the inset key. (**B**) Sequence similarity network of identified BsaA-like proteins. Nodes representing labelled *Clostridium* and *Bacillus* proteins in panel A are labeled with an arrow. Nodes are colored by taxonomy according to the color key. Proteins with conserved residues that would be expected to be involved in formation of a spontaneous intermolecular isopeptide bond are represented by triangle nodes with a red outline. (**C**) Example gene neighborhoods of genes encoding BsaA-like proteins and genes encoding putative surface filament structures. Hatching indicates the presence of the conserved NEK triad in the encoded protein. The complete matrix tree data set is included in a Supplemental Information file.

In addition to BsaA, we identified three other uncharacterized proteins in *C. perfringens* (BsaB, CPE2262 and CPE0554), but these paralogs are not expected to form an intermolecular isopeptide bond, because the N-E-K triad is missing. Nevertheless, over 100 BsaA-like proteins in other species were identified with the N-E-K triad, suggesting conservation of the donor-strand-exchange-and-lock mechanism. These proteins are largely from Clostridia with some homologs from *Coprobacillaceae*, *Enterococcaceae*, and *Thermococcus* (Fig. 9).

Based on the putative functions of gene neighbors, the BsaA/TasA family members appear to be involved in the assembly of surface filament structures (Fig. 9C). They are often in putative operons with genes encoding proteins containing the Gram-positive cell wall anchor motif LPxTG, fibronectin domains, SpaA domains, choice-of-anchor A domains, as well as signal peptidase and sortase domains. Multiple BsaA/TasA paralogs can be found in a single putative operon (Fig. 9C), suggesting the possibility of donor-strand exchange assembly of heteropolymers.

## Discussion

The original goal of this work to determine if Gram-positive bacteria have the functional equivalent of TTSS that are found in Gram-negative bacteria. Since TTSS have a similar overall architecture and mechanism as T4P (Fig. 1), it made sense to test a Gram-positive bacterium known to have T4P. With a few notable exceptions (e.g., *Streptococcus sanguinis* (29–31)) amongst Gram-positive bacteria, T4P are confined within the Clostridia (9) so *C. perfringens* was chosen as representing a typical clostridial species. The overall concept was that the thick Gram-positive PG layer represents a similar barrier to protein secretion to the environment as the Gram-negative outer membrane. Since TTSS depend upon a pseudopilus to function, we tested mutants with deletions in each of the four main pilin-encoding genes in *C. perfringens* for changes in the secretome compared to a WT strain. We identified one protein BsaC (CPE0517) that was depleted in the secretome of a *pilA3* deletion strain (Fig. 3A-B). We then examined a bank of in-frame deletion mutants in all of pilin-associated genes (Fig. 2) and found that, for BsaC secretion, a homolog of each of the essential proteins needed for a type IV pilus assembly, PilA3 (pilin), PilB2 (assembly ATPase), PilC1 (inner membrane core protein), PilD (prepilin peptidase), and PilM-N-O, (inner membrane accessory proteins) (Fig. 2) were required for efficient secretion (Fig. 4). This appears to be specific for BsaC secretion, since the secretion of the toxin PLC was not affected by mutations in any of the T4P-associated genes (Fig. S2). While this may indicate the presence of a TTSS like apparatus in *C. perfringens*, the lines separating T4P- and T2SS-dependent secretion have been blurred by the discoveries that *Pseudomonas aeruginosa* and *Dichelobacter nodosus*, Gram-negative bacteria, have T4P systems capable of mediating protein secretion (32, 33). However, further analysis of the structure and components of these Gram-negative pilus-dependent secretion systems have not been published. Since PilA3 pilin was detected on the surface of *C. perfringens* (Fig. S1), it is possible the PilA3-dependent secretion system acts more like a T4P.

BsaC monomer secretion may appear to be independent of BsaA oligomers, but it is possible that they are being secreted simultaneously, since we did not track BsaA and BsaC secretion at the same time. The biological role that BsaC plays in biofilm matrix functions is unknown, but it is intriguing that the protein contains a VWA domain, which is widespread in eukaryotes, bacteria and archaea and is associated with protein-protein interactions (34). Therefore, the role BsaC plays in biofilm functions is also currently under investigation in the laboratory.

As noted above, BsaC is in an operon with genes encoding a signal peptidase, SipW and two proteins (BsaA and BsaB) with homology to the main biofilm protein of *B. subtilis*, TasA (Fig. 3C). Because this gene synteny is similar to the *sipW-tasA-tasB* biofilm operon in *B. subtilis*, we expressed the entire operon to see if a biofilm matrix was produced and noted that high levels of a gel like matrix material was formed in the culture supernatant (Fig. S3A-B). A high mol wt oligomer of BsaA was seen in SDS-PAGE gels (Fig. S3D), which was similar to that seen by Obana et al in their analysis of the same operon (11). Because BsaC was dependent on T4P-associated proteins for secretion, it was logical to determine if secretion of the BsaA oligomer was as well. We used densitometry of western blots of the BsaA oligomers in culture supernatants that had been concentrated by TCA precipitation. Although we saw significantly decreased levels of secretion in several T4P mutants, it did not give conclusive evidence that a full T4P system was needed for secretion due to the absence of a PilC or PilD component. This experimental approach was hindered by relatively high amounts of experimental variability in trying to measure an oligomer in solution; this can be seen by the high range of the SEM determined for most of the samples (Fig. 4) despite multiple repetitions of each experiments. However, the fact that several genes in each type of experiment was shown to be necessary for efficient secretion does strongly suggest that T4P proteins play a role in the process.

Although we observed that a *pilA1* deletion strain did not secrete the biofilm matrix oligomers efficiently (Fig. 5), we did not observe this phenotype in our initial screening for proteins absent in the secretome, as shown for PilA3 in Fig. 3A-B. We think this is likely due to the fact that our growth conditions for the experiment, overnight culture in BHI liquid medium at 37 °C, is not optimum for production of the biofilm matrix proteins, as described by Obana et al (11). These findings point out the limitations of screening for T4P-dependent secretion: using only a single type of environmental condition may not be the one in which a protein is made and secreted so it would not be discovered. Even 2-D gels would be unlikely to have uncovered the presence of the BsaA oligomers due to their extremely high mol weight. Therefore, it seems protein-based mass spectrometry on samples from a wide variety of environmental conditions would be optimal for detecting T4P-dependent secretion substrates in future experiments.

Since our results and that of Obana et al both showed that oligomers of BsaA were extremely resistant to heat and other denaturants (Fig, S3C and (11)), this suggested that covalent bonding might be involved. Subsequent analysis using AlphaFold 3 indicated there were likely isopeptide bonds being formed by donor strand exchange involving an N35 of one monomer and a K74 of another, catalyzed by an E56 residue in the recipient monomer (Fig. 6). This model was confirmed by subsequent substitution of each residue involved with an Ala, which led to a complete lack of oligomerization (Fig. 7). Isopeptide bonds are formed relatively slowly (23) so we hypothesized that the donor strand exchange resulted in formation of a temporally stable interaction that gave the isopeptide bonds time to form. However, based on experiments with the N35A mutant, the oligomers are stable even without the isopeptide bond, as the mutant version was resistant to denaturation in SDS-PAGE buffer and being run through an SDS-PAGE gel. Only after heating in the SDS-PAGE buffer at 95 °C did the N35 mutant depolymerize (Fig. S4), while the WT oligomer withstood this treatment. We further characterized the formation of the oligomer and showed, using FAST fluorescent protein fusions, that the oligomers were anchored to the surface of the bacteria (Fig. 8) but via the cytoplasmic membrane and not directly to the PG layer (Fig. S5).

BsaB, the other biofilm protein encoded in the operon, did not form oligomers, did not become covalently linked to BsaA oligomers (Fig. S6) and both BsaA and BsaB have no apparent LPXTG motif present in their protein sequence (data not shown). These structural features seem somewhat at odds with the presence of a sortase gene (*srtB*) adjacent to, but oriented in the opposite direction, of the *sipW* gene (Fig. 3C). Obana et al (11) tested the premise that SrtB was responsible for oligomerization of BsaA, but found that a *srtB* mutant had normal levels of oligomer present (11). However, examination of the *bsa* operon in other strains of *C. perfringens* reveals that a different gene synteny is in place, where *sipW* is followed by a *bsaA* homolog and two *bsaB*-like genes, the first of which encodes an identifiable LPXTG motif (Fig. 3D). The BsaB sequence from strain 13, when subjected to a BLAST analysis (35) against other *C. perfringens* strains, shows proteins with high levels of homology to either the N-terminal half or C-terminal half of the strain 13 BsaB, but not both. This suggests strain 13 underwent a genetic rearrangement event in which the two *bsaB* homologs were recombined into a single gene with the loss of the LPXTG motif found in the first gene (Fig. 3C-D). This brings up the distinct possibility that the SrtB sortase could catalyze the cross-linking of the first BsaB ortholog to the PG layer in strains other than strain 13. What role BsaB plays in BsaA oligomerization is still unknown, since a *bsaB* mutant still formed similar amounts of oligomers of BsaA as the wild-type strain (11). However, based on the differences in composition of BsaB homologs between strain 13 and other strains (Fig. 3C-D), it is possible that BsaB is involved in anchoring the BsaA oligomers to the PG layer via the activity of SrtB and the BsaB homolog with an LPXTG motif, but this is speculative.

Taking all these results into consideration, we have constructed a model showing the mechanisms underlying the assembly of BsaA into cross-linked oligomers (Fig. 10). In this model, BsaA monomers are secreted via the Sec pathway, fold on the outer surface of the CM but are anchored in the membrane by their N-terminal signal sequence. An oligomer of BsaA with the most recent monomer having its N-terminal disordered region exposed, contacts the newly secreted monomer and undergoes a donor strand exchange in which the N-terminal disordered region binds to the exposed beta-sheet fold and interdigitates as a newly formed beta sheet (Fig. 6A). This is followed by cleavage of the signal sequence by the signal peptidase SipW. We believe the SipW acts after oligomer binding to the newly secreted monomer because the BsaA oligomers appear to be anchored to the membrane (Fig. S5) and the oligomers are in a relatively stable assembly even before the monomers are “locked” together by isopeptide bond formation (Fig. S4). After SipW cleavage, the oligomer is released and can then be used to add another monomer or be secreted through the PG layer to the external medium. The presence of significant amounts of oligomers attached to the bacterial surface (Fig. 8) suggests the monomers assemble in the space between the CM and PG rather than being secreted as monomers for oligomerization in the environment. The mechanism for export of the oligomers through the PG layer is not complete but we provided evidence that T4P play at least some role in the process (Fig. 5).

**Figure 10.**
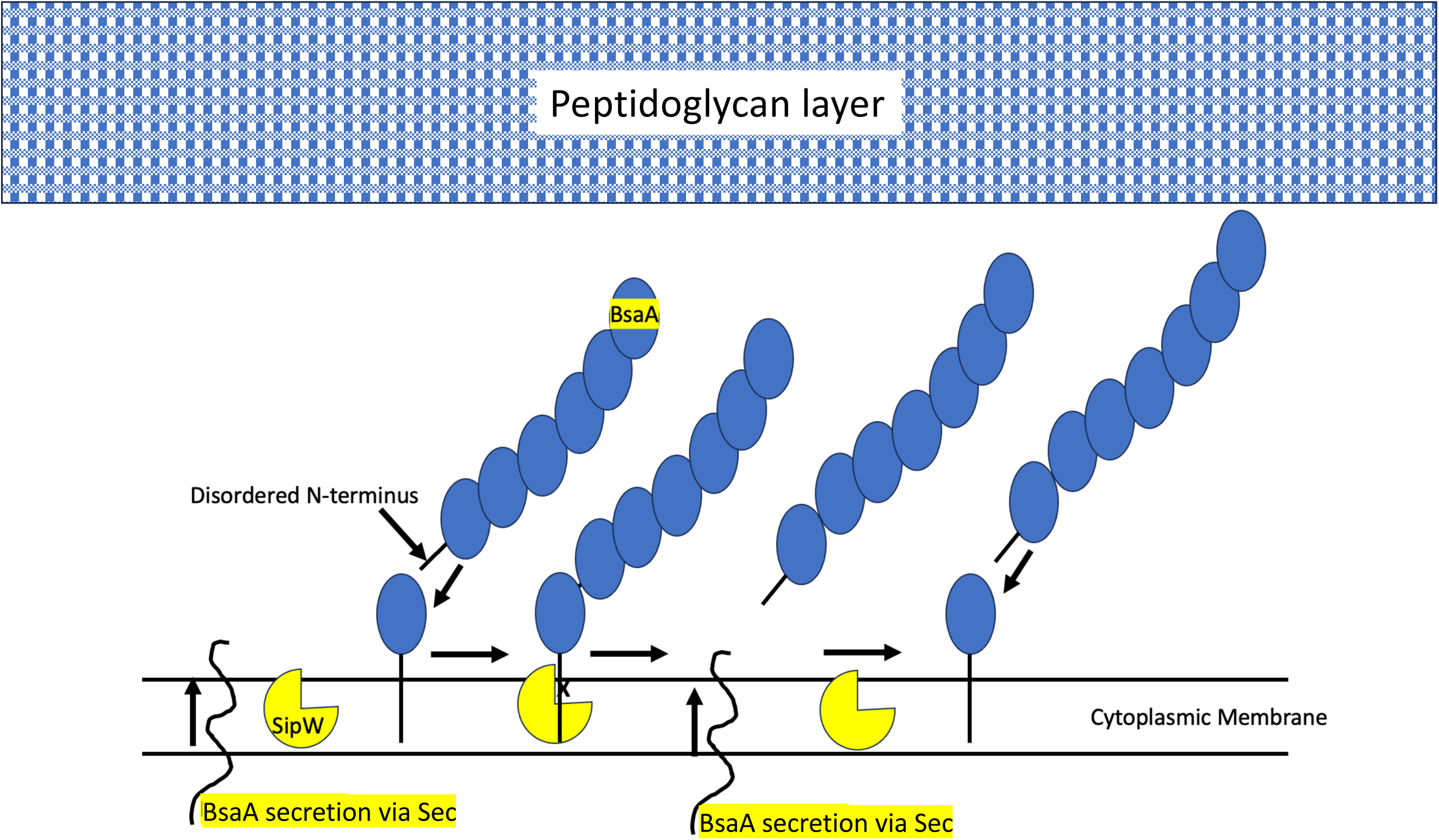
Model of BsaA oligomerization. BsaA monomers (purple ovals) are oligomerized by isopeptide bond formation after secretion via the Sec system in *C. perfringens*. SipW is the signal peptidase for BsaA secretion (Fig. 3C and (11)). Secretion of the oligomers past the PG layer may be due to the action of T4P-associated proteins (Fig. 5).

The phylogeny of BsaA-like proteins shows they are widespread amongst Gram-positive bacteria but also present in some Gram-negative bacteria and archaea (Fig. 9). However, the homologs that are predicted to form isopeptide bonds are more narrowly focused in the Clostridia and Enterococcus genera (Fig. 9). This brings up the question as to what advantage isopeptide bond formation bestows on the biofilm matrix properties of the bacteria that form them. Formation of isopeptide bonds can greatly increase the mechanical resistance to distension and separation of the BsaA fibers. As an example, the intramolecular isopeptide bonds formed in sortase-dependent pili made by Gram-positive bacteria have been shown to greatly increase the amount of force that needs to be applied to distend the individual subunits, thereby strengthening the entire fiber (36). To trap macrophages, neutrophils or other phagocytic cells, *C. perfringens* and other species may have evolved a mechanism to form isopeptide bonds to increase the mechanical strength of the fibers, since macrophages, neutrophils and other phagocytic cells can impart a large locomotive force (1.9 to 10.7 nN) during their motility (37). It is also worth noting that *C. perfringens* is an extremely ubiquitous bacterium in nature, found in soil, freshwater sediments and even extreme environments like Antarctica (38, 39) and likely forms biofilms in these natural environments. The BsaA biofilm matrix fibers could protect them from killing by amoeba, protozoa and other predators found in these environments. Analyzing the relative mechanical strength of fibers with and without isopeptide bonds will help answer some of these questions.

## Materials and Methods

### Bacterial strains and growth conditions

Bacterial strains, plasmids, and primers used in this study are listed in Table 1. *Escherichia coli* strain DH10B was grown in Luria Bertani broth at 37°C for all transformations. When necessary, kanamycin and chloramphenicol were added to the media at a concentration of 100 μg/ml and 20 μg/ml, respectively. *C. perfringens* strain HN13, a Δ*galKT* derivative of strain 13, was used as the wild type strain in this study. *C. perfringens* strains were grown anaerobically in PGY (30 g proteose peptone #3, 20 g glucose, 10 g yeast extract, 1 g sodium thioglycolate per liter) or brain-heart infusion (BHI) (Thermo Fisher) in an anaerobic chamber (Coy Laboratory Products, Inc.). Anaerobic egg yolk agar medium was prepared as previously described (https://www.fda.gov/food/laboratory-methods-food/bam-media-m12-anaerobic-egg-yolk-agar#).

**Table 1.**
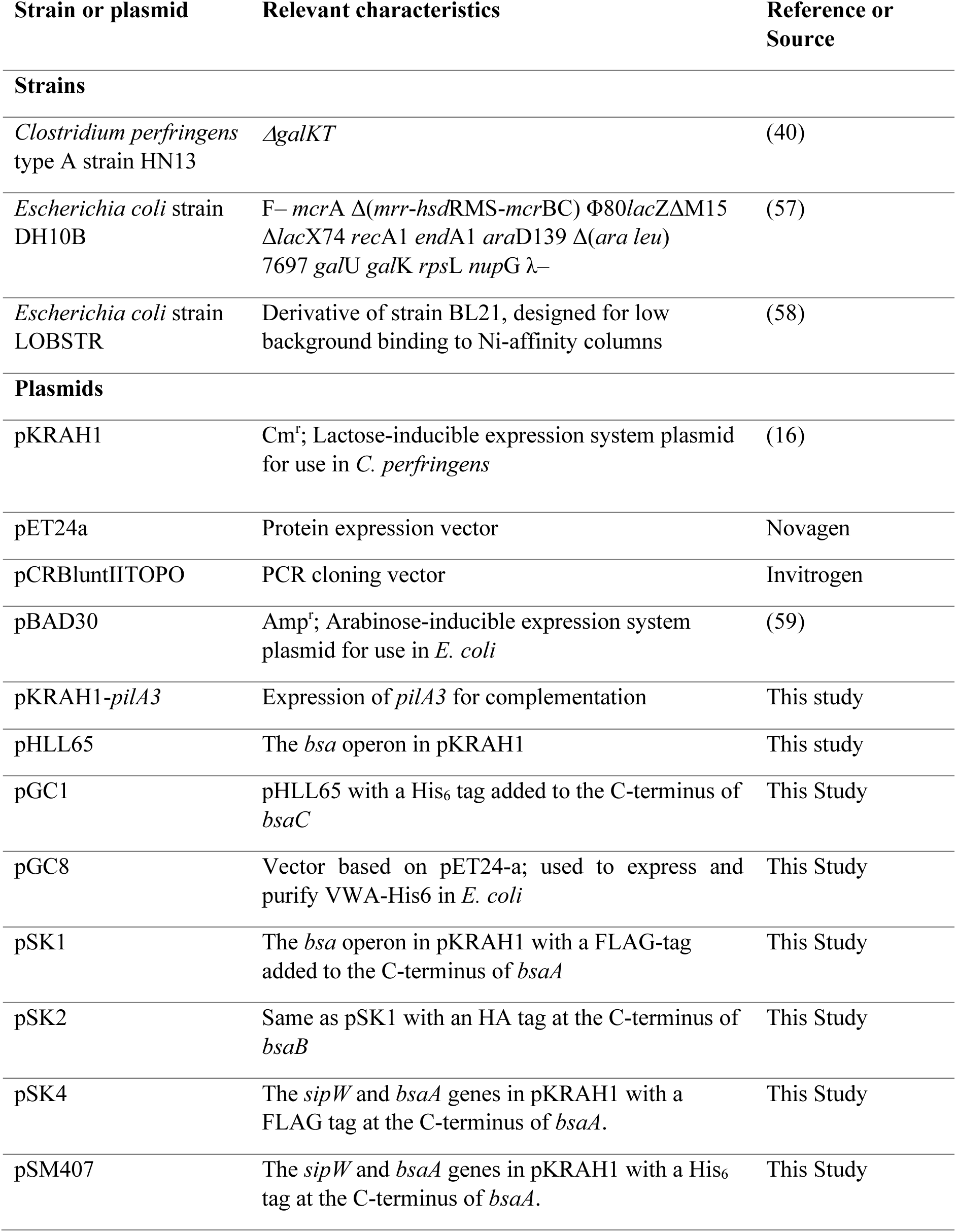
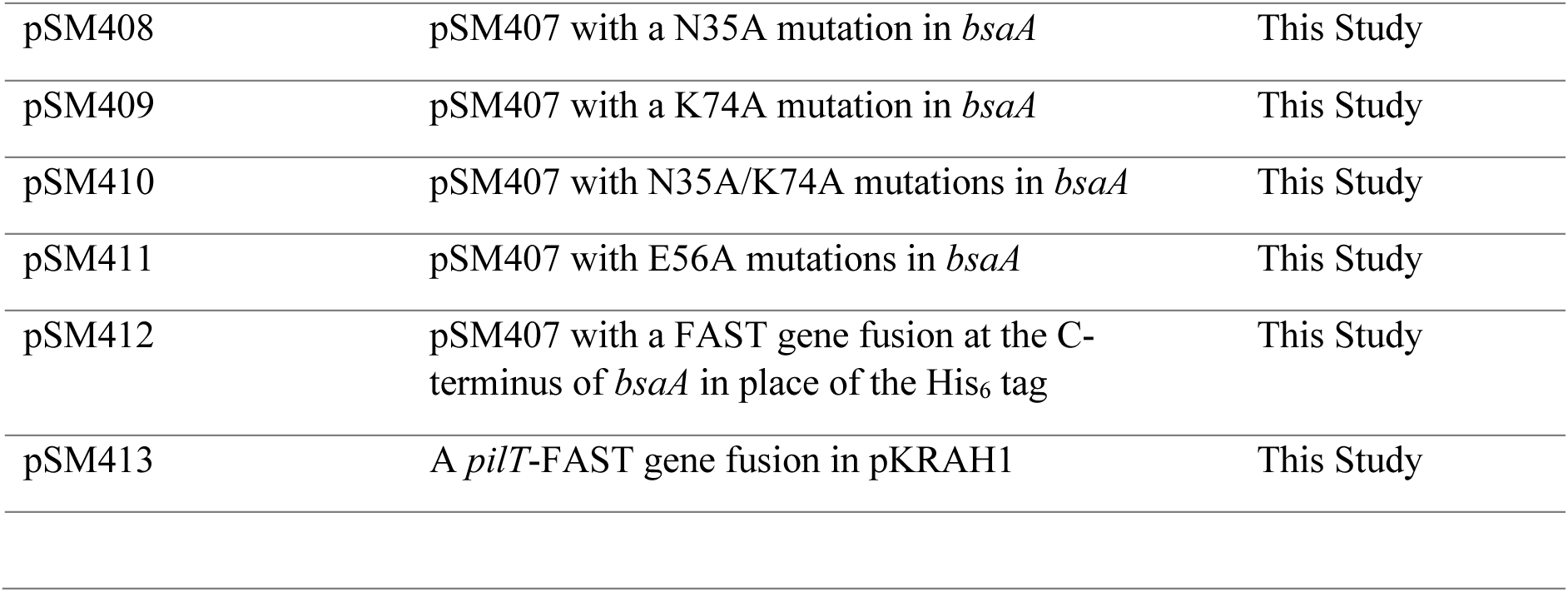
Strains and plasmids used in this report.

### Construction of in-frame deletions of T4P-associated genes

In-frame deletions of T4P-associated genes were made using the method of Nariya et al. (40), modified as described in Hendrick et al (41). The primers used to amplify the flanking DNA for each gene are listed in Table S1. All deletions were confirmed by PCR across the deleted region. Complementation of the in-frame *pilA3* mutant (Fig. 3A-B) was done using the lactose-inducible promoter vector pKRAH1 (16) to express the *pilA3* gene. The construct was made by PCR using the primers listed in Table S7.

### SDS-PAGE and staining of secretome proteins

Cultures of strain HN13 were grown for 16 h in BHI medium (i.e., into stationary phase) and the bacteria were removed by centrifugation in a table top centrifuge. The culture supernatants were removed and passed through a 0.45 μM filter. Proteins in the culture medium were precipitated using 10% trichloroacetic acid (TCA) precipitation and washed twice in acetone. After drying, sample pellets were resuspended in water and then 4X SDS-PAGE sample buffer (42) was added to a final concentration of 1X. Secretome samples were separated on an SDS-PAGE gel and then stained with the fluorescent protein stain, Sypro-Ruby, according to the manufacturer’s instructions (Thermo Fisher). Stained gels were imaged with a Gel-Doc Imaging system (Bio-Rad, Inc).

### Cloning the *bsa* operon into the lactose-inducible expression vector, pKRAH1

Because the SipW signal peptidase is likely needed for processing the secretion signal on BsaC (11), to analyze secretion of BsaC in T4P gene associated mutants, the entire *sipW-basA-bsaB-bcaC* operon was expressed under control of the lactose inducible promoter in plasmid pKRAH1 (16) using primers OHL145 (PstI site) and OHL146 (BamHI site) by PCR (Table 1 and Table S3). The PCR fragment was ligated to the PCR cloning vector pGEM-T to make pHLL64. pHLL64 and pKRAH1 were digested with PstI and BamHI and ligated to make pHLL65.

### Construction of a FAST protein fusion to BsaA

The gene fusion was constructed using overlapping PCR of the FAST coding region on plasmid pAG104 FAST (26) to the *bsaA* gene linked to *sipW* (i.e., the *bsaB* and *bsaC* genes were absent). The FAST protein was fused to the C-terminal end of BsaA because of the processing of the signal sequence on the N-terminal end. The primers used to construct the gene fusion are listed in Table S5.

### Production of antibodies to BsaC, PilA3 and immunofluorescence microscopy

Rabbit polyclonal antibodies against the BsaC peptide sequence N-CNIEGDKISWDRGEN-C were produced by GenScript USA, Inc. along with pre-immune serum as a negative control. Rabbit polyclonal antibodies against the PilA3 peptide sequence N-CIKIDNSKDSIDLTR-C were produced by Yenzyme Antibodies, LLC. along with pre-immune serum as a negative control. To test for surface exposure of PilA3 pilins, the bacteria were fixed with 2.5% paraformaldehyde in Dulbecco’s phosphate buffer saline (DPBS), incubated with rabbit serum with anti-PilA3 antibodies, washed with DPBS and incubated with goat anti-rabbit antibodies fluorescently tagged with Alexafluor 647 (Invitrogen). Stained cells were imaged on an Olympus IX71 fluorescent inverted microscope equipped with a CoolSnap HQ2 CCD camera and DeltaVision deconvolution and image analysis software.

### Quantitative immunoblotting of secreted BsaC

Overnight cultures were grown in BHI supplemented with chloramphenicol and diluted 1:50 into fresh BHI medium pre-incubated at 37°C. The cultures were grown for one hour then induced by adding lactose, to a final concentration of 10 mM, to the growth media every 30 min for 3 h. Following induction, 1 mL of culture supernatant was collected by centrifugation, and proteins precipitated using TCA, as described above. Resulting precipitates were resuspended in 50 uL of SDS-PAGE loading buffer with 100 mM DTT, the pH was adjusted using one μL of 1.5 M Tris-HCl, pH 9, if necessary. Five μL of sample was loaded per well, and SDS-PAGE was performed. Following SDS-PAGE, proteins were transferred to a polyvinylidene difluoride (PVDF) membrane utilizing the iBlot system (Bio-Rad, Inc), and developed with serum containing anti-BsaC antibodies at 1:250 dilution, followed by a Dylight 550 labeled fluorescent secondary goat anti-rabbit antibody (Thermo-Fisher). Imaging was performed on a Typhoon gel scanner, and mean pixel intensity was measured for each band utilizing the Amersham Typhoon densitometry control software.

### Purification of BsaC

The *bsaC* gene with a C-terminal His_6_ tag was cloned into the expression vector pET24a by PCR using the primers listed in Table S4. An overnight culture of *E. coli* LOBSTR pET24a::BsaC-His_6_ was used to inoculate 2 L of 37°C pre-warmed terrific broth in a 4 L beveled flask. The culture was allowed to grow to 0.6 OD_600_ where it was induced with 1 mM IPTG. After 6 hours of induction, the culture was centrifuged to remove cells (BsaC-His_6_ is secreted by *E. coli*), and the resulting supernatant was precipitated with 90% ammonium sulfate. Precipitated protein was solubilized in PBS and used to load on to an ÄKTA purifier FPLC for Ni-NTA affinity chromatography in PBS with 10 mM imidazole as loading buffer, and PBS with 1M imidazole as elution buffer. Eluted fractions were concentrated with an Amicon Ultra 2 mL 10 K cutoff centrifugation columns. Concentrated elution fractions were then further processed using a Superose 6 gel filtration column linked to an ÄKTA purifier FPLC.

### Immunoblotting of BsaA oligomers

BsaA oligomer secretion by *C. perfringens* was analyzed using western blots. For the immunoblot experiments, the proteins were blotted onto a PVDF membrane while submerged in cold buffer which contained 3 mM Na_2_CO_3_, 10 mM NaHCO_3_ and 20% methanol.

### Immunoblotting Method One

The membranes were placed in a Snap i.d. 2.0 system (Millipore Sigma) which uses a vacuum system to pull solutions through the membrane (referred to as washing steps below). Once set up, a solution containing Tris-buffered saline (TBS) (pH 7.4), 0.1% Tween 20 (TBST), and 1% bovine serum albumin (BSA) was added as a blocker. Both primary and secondary antibodies were added to separate aliquots of the solution containing TBST and 1% BSA. In the first step, the primary antibody solution was incubated for 10 minutes on the membrane, followed by three washings with TBST. Then the secondary antibody solution was left to sit for 10 minutes on the membrane, followed by three washings with TBST. The primary antibody utilized was OctA-probe H-5 (1:250 dilution) used to detect the FLAG epitope (Santa Cruz Biotechnology). The secondary antibody utilized for these experiments was StarBright700 goat anti-mouse (Bio-Rad) (1:10,000 dilution). The fluorescently labeled immunoblots were imaged using a ChemiDoc MP imaging system (Bio-Rad), and the intensity of the fluorescence was measured using the ChemiDoc MP imaging software for densitometry measurements.

### Immunoblotting method 2

Membranes were transferred to a container and blocked with EveryBlot blocking Buffer (Bio-Rad). Antibodies were diluted with EveryBlot blocking buffer and applied to membranes, followed by a minimum of five washes with TBST.

The primary antibodies utilized included OctA-probe H-5 (1:227 dilution) used to detect FLAG, and HA-Tag F-7 (1:227 dilution) used to detect HA (both from Santa Cruz Biotechnology, Santa Cruz, CA). Membranes submerged in EveryBlot blocking buffer with primary antibody were incubated at 4°C overnight. The secondary antibody utilized for these experiments was StarBright700 goat anti-mouse (Bio-Rad) (1:13,333 dilution), incubated for one hour at room temperature. Images of the immunoblots were taken using a ChemiDoc MP imaging system (Bio-Rad), to measure the intensity of the fluorescence.

### Western blots on *bsaA* mutants

Western blots to detect oligomerization of BsaA mutants were performed by expressing the *sipW-bsaA-His_6_* genes under control of the lactose-inducible promoter. Supernatants were separated from cell pellets by centrifugation. The proteins in the supernatant were concentrated by TCA precipitation (as described above), suspended in SDS-PAGE loading buffer with 100 mM DTT and heated at 95 °C for 20 min. The bacteria in the cell pellet were suspended in PBS and disrupted by bead beating with 0.1 mm zirconia-silica beads. The beads were then removed by centrifugation and the cell extract brought to 1 X SDS-PAGE buffer with 100 mM DTT and heated at 95 °C for 20 min. After running on SDS-PAGE gels, the proteins were transferred to a PVDF membrane and developed using immunoblotting method 2 above, except the primary antibody was mouse monoclonal anti-His6 antibody His.H8 (Santa Cruz Biotechnology).

### Modeling BsaA Structure

BsaA from *C. perfringens* (accession number WP_011009871) was created as a homotrimer using ColabFold v1.3.0 (19) based on AlphaFold v2.2.0 (20). Amber minimization was not enabled. In the top ranked model based on pLDDT PyMOL v2.5.0 was used to analyze side chains of interdigitated beta strands for potential sidechain crosslinks. Images were created using ChimeraX v1.2. (43, 44) The same trimer of BsaA was created in AlphaFold 3 (25) to observe if the new version creates crosslinks, and all trimer models made did contain the crosslinks. Structures of BsaA were created with AlphaFold 3 (25).

### Phylogeny of BsaA-like proteins

BsaA-like protein sequences were identified with 3 iterations of jackhmmer (45) (HmmerWeb version 2.41.2) (46) using BsaA (UniProt:Q8XN23) from *C. perfringens* as the seed. The protein sequence similarity network of proteins identified either with jackhmmer or as a match to IPR022121 was constructed using the EFI-EST tool (https://efi.igb.illinois.edu/efi-est/) (47, 48) with an alignment score of 10. Nodes were collapsed at a sequence identity of 100%. The network was visualized with Cytoscape (https://www.cytoscape.org) (49) using the Prefuse Force Directed OpenCL Layout. Gene neighborhoods were retrieved using the EFI-GNT tool (https://efi.igb.illinois.edu/efi-gnt/) (48). The presence of a N-E-K triad was determined by manual inspection of the multiple sequence alignment. For phylogenetic reconstruction, sequences were aligned with MAFFT on XSEDE using CIPRES (50, 51), edited in Jalview (52), and analyzed with IQ-Tree (53–55). Maximum likelihood trees were constructed using the LG+F+G4 substitution model and ultrafast bootstrap (1000 replicates). Consensus trees were visualized, rooted at the midpoint, and annotated with iTOL (56). Data files are available in Supplemental File Data Set 1.

## Supporting information

Biofim matrix protein SI Data Set

## Acknowledgments

We thank W. Keith Ray, Richard Helm and the Virginia Tech Proteomics Incubator for assistance with mass spectrometry and Megan Rochford and Hualan Liu for construction of in-frame deletions mutants. Research reported in this publication was supported by the National Institute of Allergy and Infectious Disease of the National Institutes of Health under award number R21AI109391 and R21AI149177 to S. B. M. The content is solely the responsibility of the authors and does not necessarily represent the official views of the National Institutes of Health. Work at the Molecular Foundry was supported by the Office of Science, Office of Basic Energy Sciences, of the U.S. Department of Energy under Contract No. DE-AC02-05CH11231. The work conducted by the U.S. Department of Energy Joint Genome Institute (https://ror.org/04xm1d337), a DOE Office of Science User Facility, is supported by the Office of Science of the U.S. Department of Energy operated under Contract No. DE-AC02-05CH11231.

## Author Contributions

Sarah E. Kivimaki: Investigation, Methodology, Visualization, and Writing – Review and Editing.

Samantha Dempsey: Investigation, Methodology, Visualization, and Writing.

Gary J. Camper: Investigation, Methodology, Visualization, and Writing.

Julia M. Tani: Investigation, Methodology, Visualization.

Ian K. Hicklin: Investigation, Methodology, Visualization, and Writing.

Crysten E. Blaby-Haas: Methodology, Visualization, and Writing – review and editing.

Anne M. Brown: Methodology, Visualization, and Writing – review and editing.

Stephen B. Melville: Conceptualization, Funding acquisition, Supervision, Investigation, Methodology, Project administration, Writing, Review and editing.

## Notes

### Competing Interest Statement

The authors have declared no competing interest.

### Summary of Updates

The text and figures have been updated for clarity.

