## Supplemental figures for "Type IV pili-associated secretion of a biofilm matrix protein from *Clostridium perfringens* that forms intermolecular isopeptide bonds"

### Slide 1
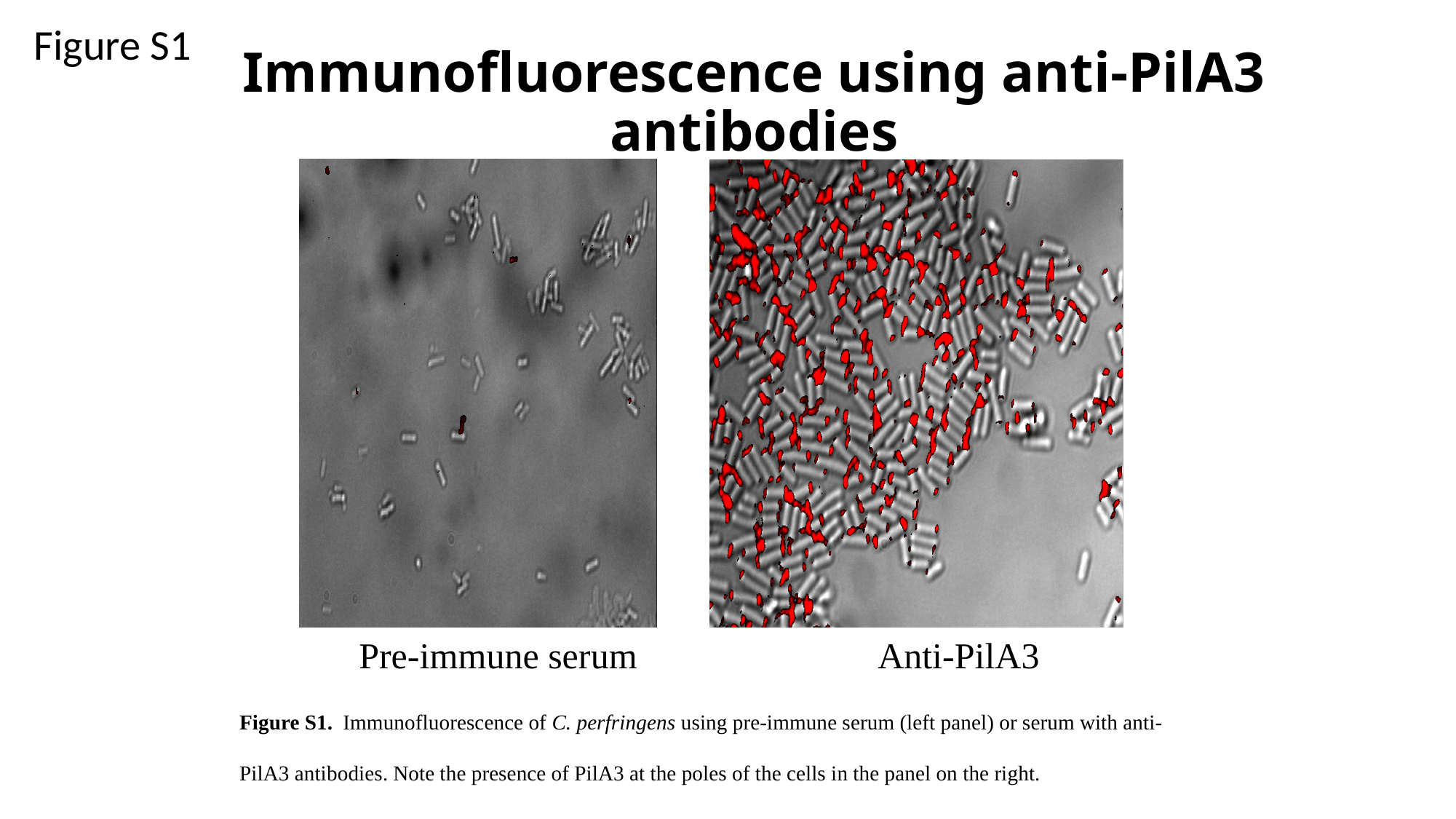

Figure S1
#### Immunofluorescence using anti-PilA3 antibodies
Pre-immune serum
Anti-PilA3
Figure S1. Immunofluorescence of C. perfringens using pre-immune serum (left panel) or serum with anti-PilA3 antibodies. Note the presence of PilA3 at the poles of the cells in the panel on the right.

### Slide 2
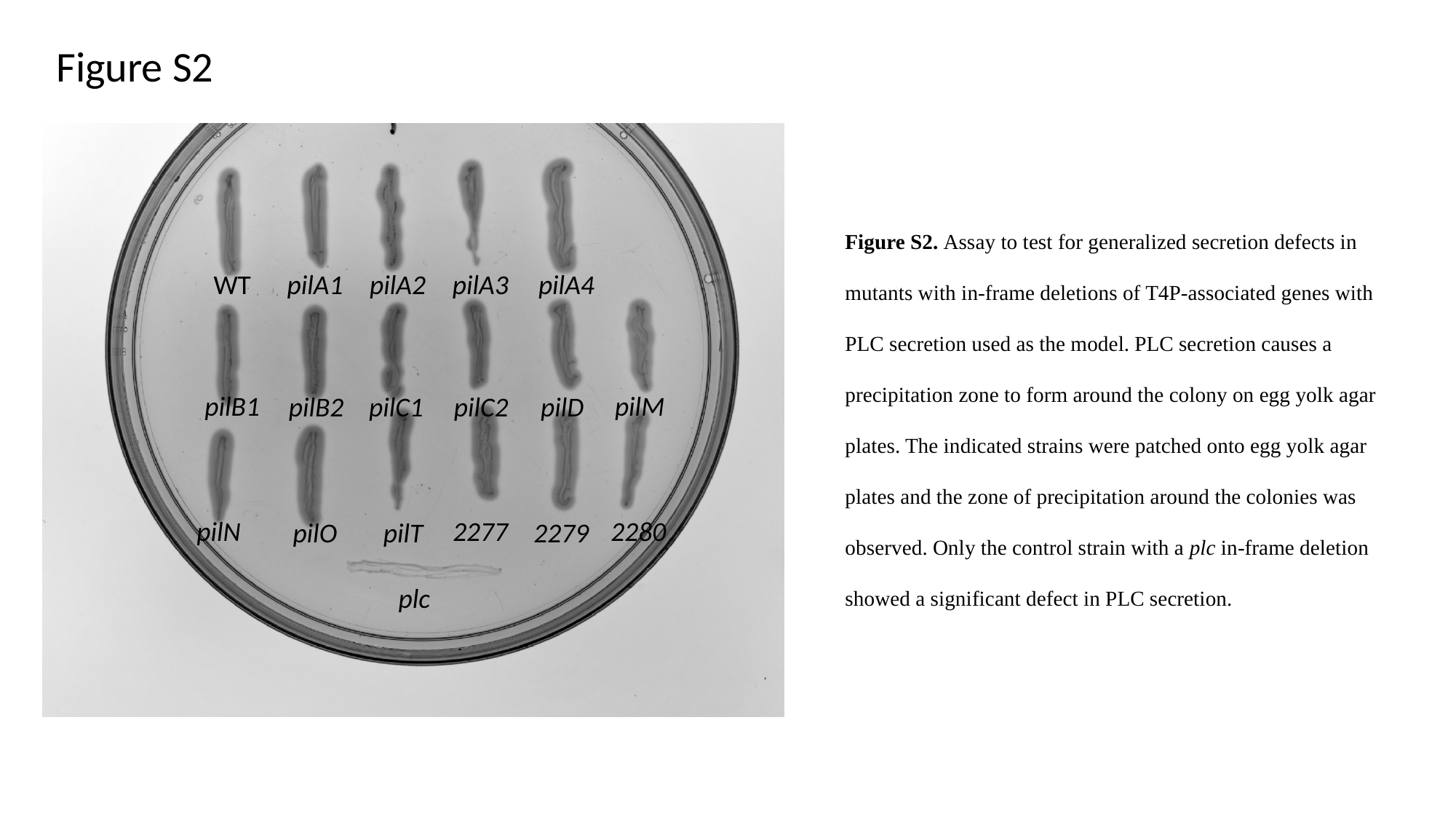

Figure S2
WT
pilA1
pilA2
pilA3
pilA4
pilM
pilB1
pilB2
pilC1
pilC2
pilD
2277
2280
pilN
pilO
pilT
2279
plc
Figure S2. Assay to test for generalized secretion defects in mutants with in-frame deletions of T4P-associated genes with PLC secretion used as the model. PLC secretion causes a precipitation zone to form around the colony on egg yolk agar plates. The indicated strains were patched onto egg yolk agar plates and the zone of precipitation around the colonies was observed. Only the control strain with a plc in-frame deletion showed a significant defect in PLC secretion.

### Slide 3
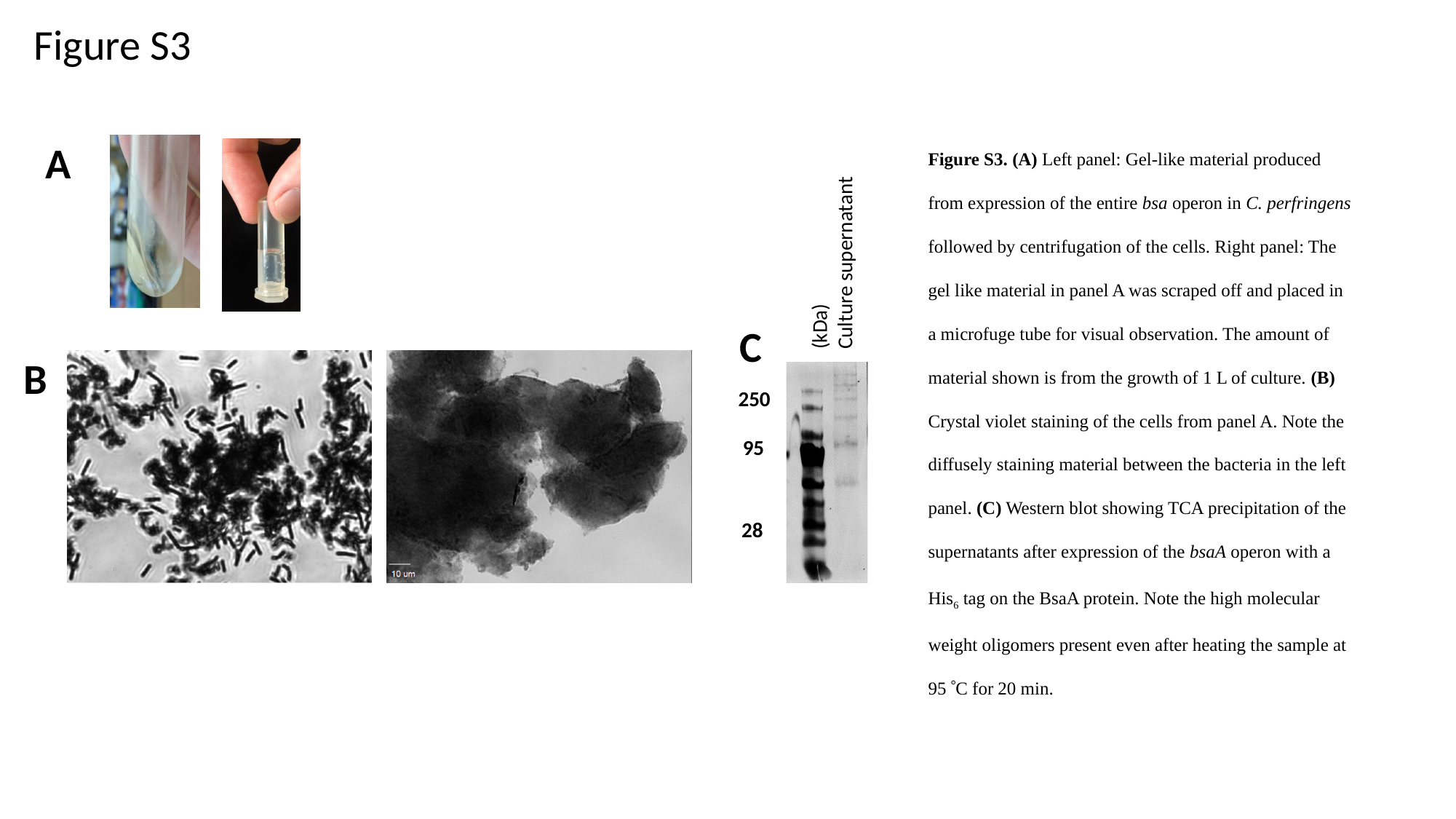

Figure S3
Figure S3. (A) Left panel: Gel-like material produced from expression of the entire bsa operon in C. perfringens followed by centrifugation of the cells. Right panel: The gel like material in panel A was scraped off and placed in a microfuge tube for visual observation. The amount of material shown is from the growth of 1 L of culture. (B) Crystal violet staining of the cells from panel A. Note the diffusely staining material between the bacteria in the left panel. (C) Western blot showing TCA precipitation of the supernatants after expression of the bsaA operon with a His_6_ tag on the BsaA protein. Note the high molecular weight oligomers present even after heating the sample at 95 °C for 20 min.
A
(kDa)
Culture supernatant
C
B
250
95
28

### Slide 4
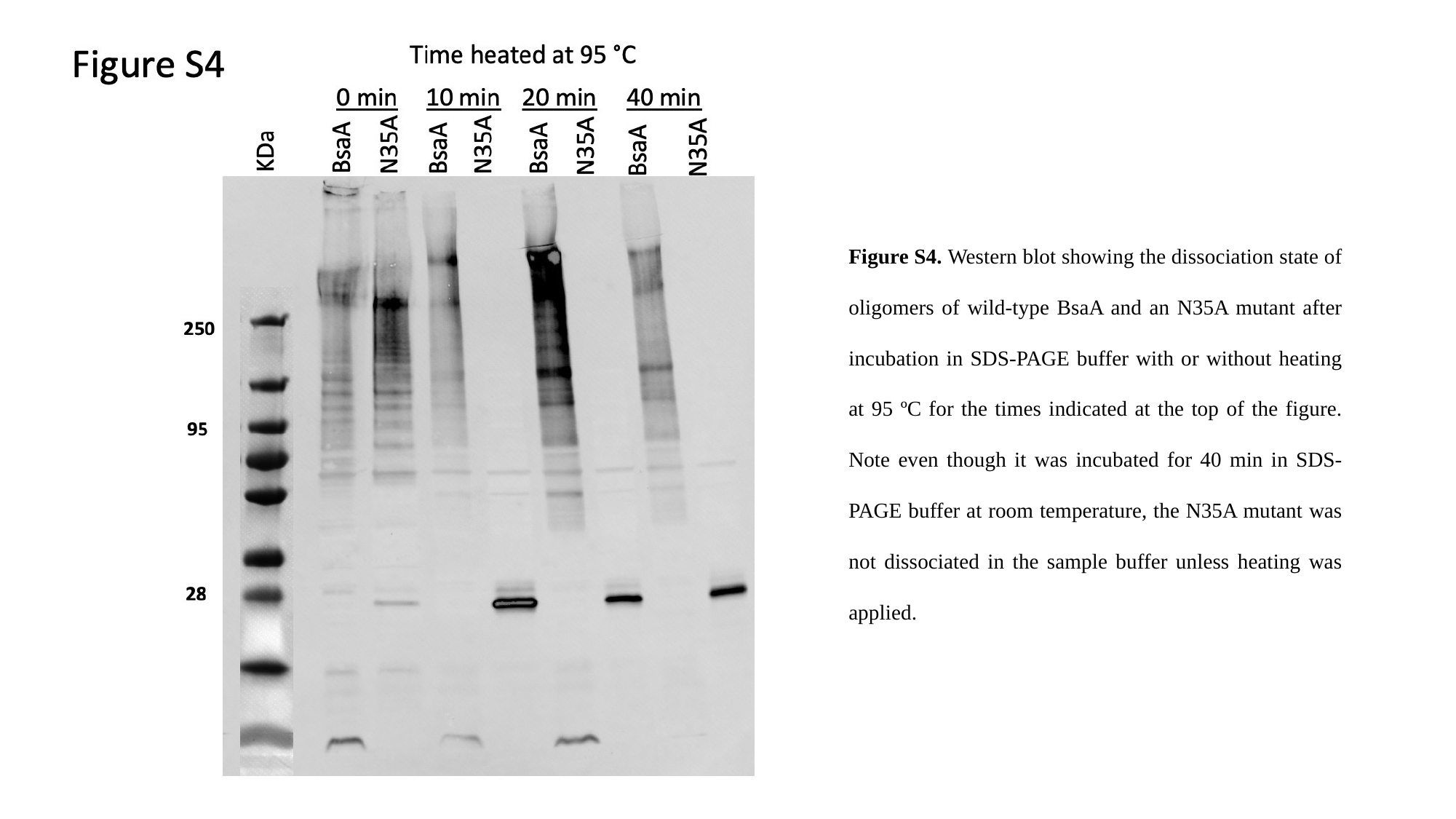

Figure S4. Western blot showing the dissociation state of oligomers of wild-type BsaA and an N35A mutant after incubation in SDS-PAGE buffer with or without heating at 95 ºC for the times indicated at the top of the figure. Note even though it was incubated for 40 min in SDS-PAGE buffer at room temperature, the N35A mutant was not dissociated in the sample buffer unless heating was applied.

### Slide 5
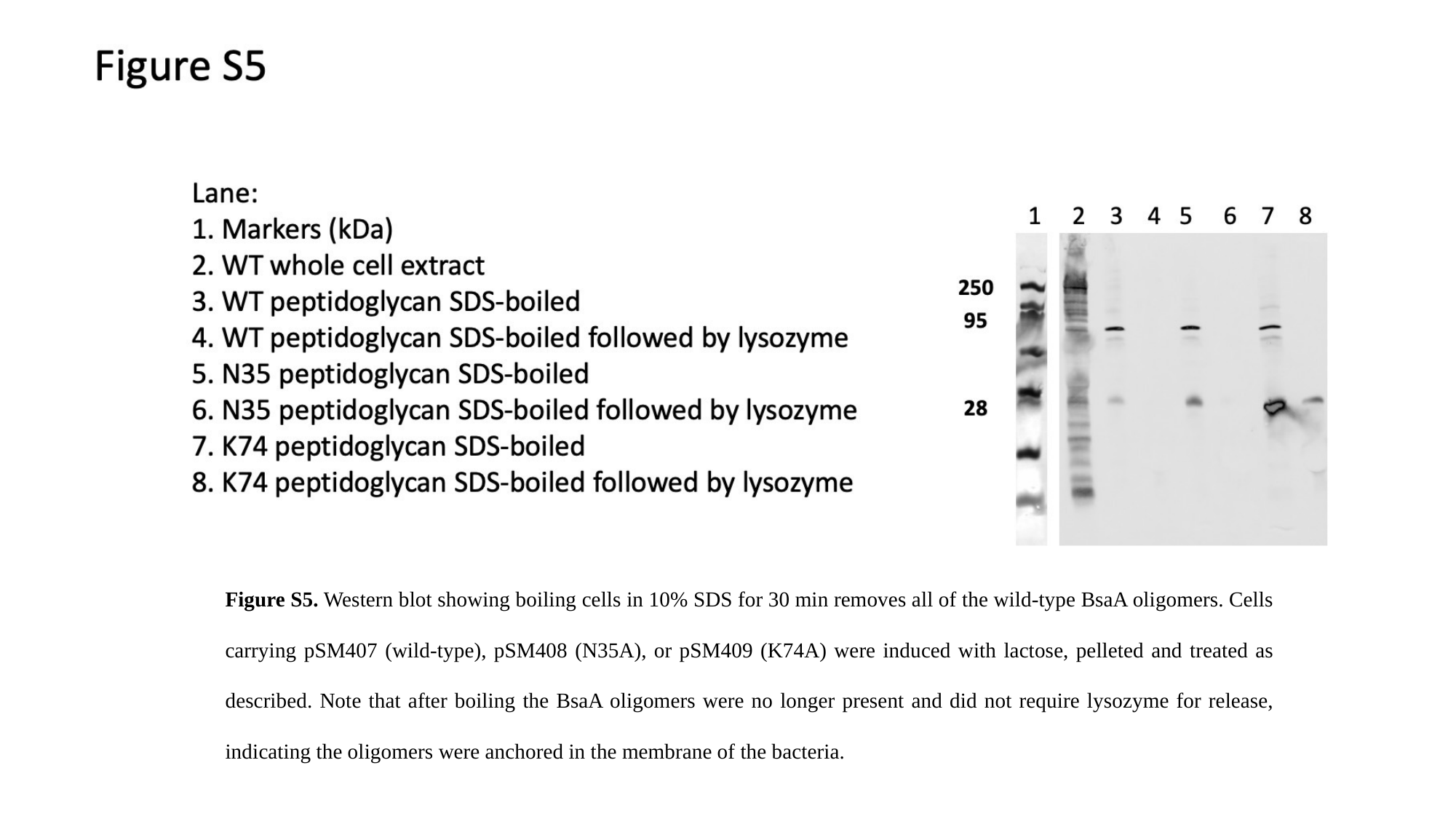

Figure S5. Western blot showing boiling cells in 10% SDS for 30 min removes all of the wild-type BsaA oligomers. Cells carrying pSM407 (wild-type), pSM408 (N35A), or pSM409 (K74A) were induced with lactose, pelleted and treated as described. Note that after boiling the BsaA oligomers were no longer present and did not require lysozyme for release, indicating the oligomers were anchored in the membrane of the bacteria.

### Slide 6
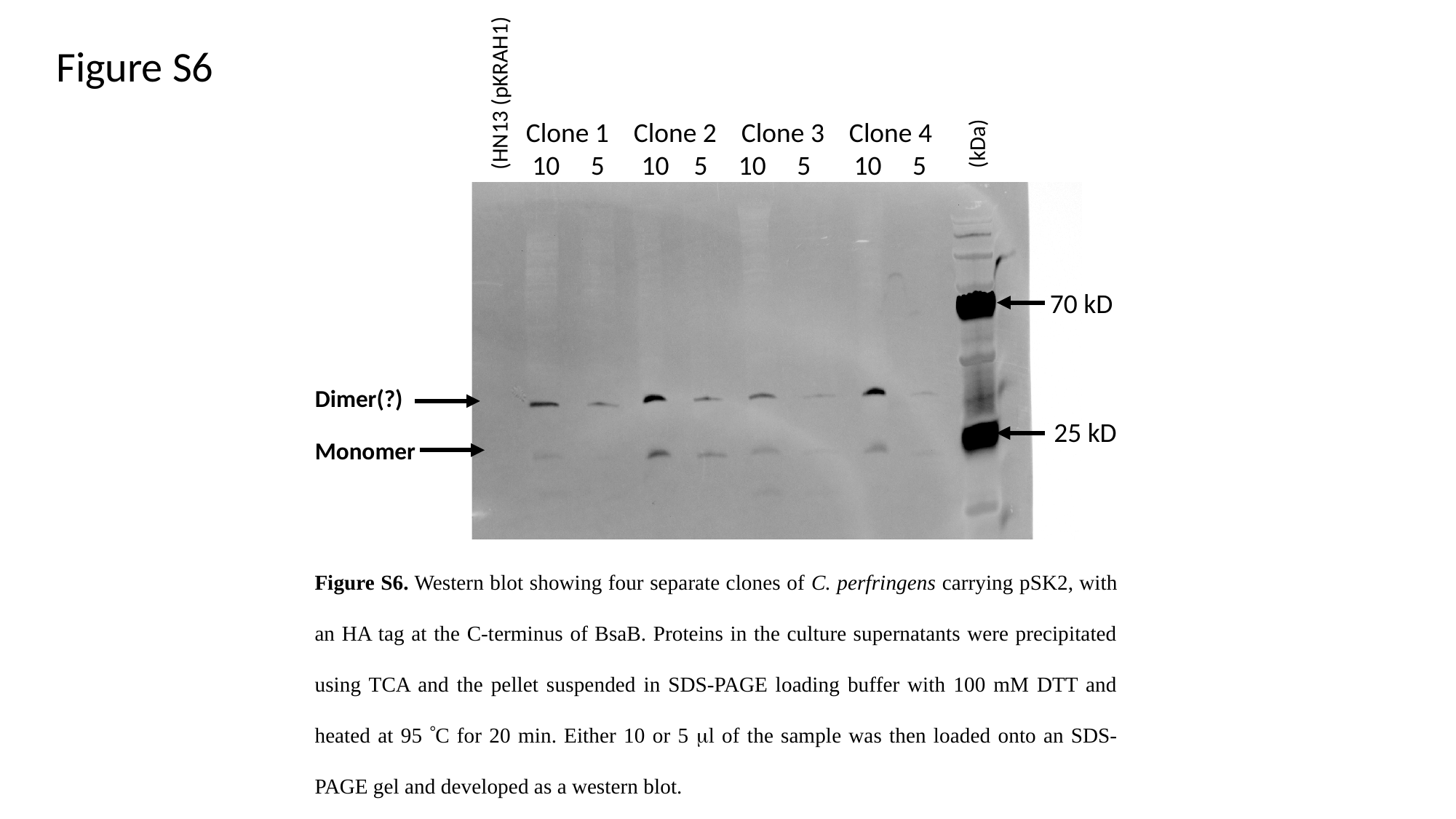

(HN13 (pKRAH1)
Dimer(?)
Monomer
Clone 1 Clone 2 Clone 3 Clone 4
 10 5 10 5 10 5 10 5
(kDa)
70 kD
25 kD
Figure S6
Figure S6. Western blot showing four separate clones of C. perfringens carrying pSK2, with an HA tag at the C-terminus of BsaB. Proteins in the culture supernatants were precipitated using TCA and the pellet suspended in SDS-PAGE loading buffer with 100 mM DTT and heated at 95 °C for 20 min. Either 10 or 5 μl of the sample was then loaded onto an SDS-PAGE gel and developed as a western blot.
