## Supplemental text and figure legends for "Type IV pili-associated secretion of a biofilm matrix protein from *Clostridium perfringens* that forms intermolecular isopeptide bonds"

**Table S1.** The primers used to construct in-frame deletions or the reference for previously published deletions. Each deletion requires four primers, the first one is the 5’ flanking primer, the next two are the overlapping mutagenic primers and the last one is the 3’ flanking primer.

| Gene deleted | Primer | Sequence (5’-3’) | Reference |
| --- | --- | --- | --- |
| *bsaA* | OSM427 | TAGGATCCTCTGGACTTTTATTAACAGAAGAATCAGATAGAGG | This report |
|  | OSM434 | CTCCTTGCATCATGGTTATAACTTAATTTAAAATATTACTTATTTATGCTTACTCAATTTAAATTACCTCCCAGTTTATAAATTATTTATATTATTC | This report |
|  | OSM435 | GAATAATATAAATAATTTATAAACTGGGAGGTAATTTAAATTGAGTAAGCATAAATAAGTAATATTTTAAATTAAGTTATAACCATGATGCAAGGAG | This report |
|  | OSM430 | CTTGTCGACGCTCTTGTGATACTTCTTTAAGAATTTCAGTTCCG | This report |
| *bsaC* | OOG1 | GGATCCACTGCAAACTTATTAGAAAGTGTTAC | This report |
|  | OOG2 | CAAAAATAAAAAGTTTTCCATCTTAATTTATCTTCATATCTCTCCCCACCTAAC | This report |
|  | OOG3 | GTTAGGTGGGGAGAGATATGAAGAATTTAAATTAAGATGGAAAACTTTTTATTTTTG | This report |
|  | OOG4 | GTCGACACATAAAATTACATATCGCCTATTC | This report |
| *sipW* | OGC16 | ATATATAGGCCTTTAATCAATTTCTTTTGC | This report |
|  | OGC17 | TTATATTATTCTTATAATTTAAATCTAACTTTCTTTTTTTTCATAATTATTCCTTTTCTTCC | This report |
|  | OGC18 | GGAAGAAAAGGAATAATTATGAAAAAAAAGAAAAGTTAGATTTAAATTATAAGAATAATATAA | This report |
|  | OGC19 | CTTTTTACTAACTCTTTCAGCTCTCAT | This report |
| *pilA1* |  |  | (1) |
| *pilA2* |  |  | (1) |
| *pilA3* |  |  | (1) |
| *pilA4* |  |  | (1) |
| *pilB1* | OAH155 | TAGCCTAGGCCATGGTCGACGGGGCATATGGAAGTTTATAGTGAG | This report |
|  | OAH156 | GTAAAACATGCCTTTAATAGTTCTTCTCCGGTATATTTCA  TATATCAACCTCCTCATCAG | This report |
|  | OAH157 | GATGAGGAGGTTGATATATGAAATATACCGGAGAAGAACT  ATTAAAGGCATGTTTTACC | This report |
|  | OAH158 | GATCTAGACTCGAGCTCCCCTATACTACTGCTTAAAGTAGTCCC | This report |
| *pilB2* |  |  | (2) |
| *pilC1* |  |  | (2) |
| *pilC2* |  |  | (2) |
| *pilD* | OSRM9 | GGAGGACCATAAATGTTATTACTGAAAGCAAGTGCAATATGTGG | This report |
|  | OSRM10 | CTAGTATATTAAATTTTTCACACACCAAATATAGACACATTGCCTCCCAAATCATCC | This report |
|  | OSRM11 | GGATGATTTGGGAGGCAATGTGTCTATATTTGGTGTGTGAAAAATTTAATATACTAG | This report |
|  | OSRM12 | CCACTCCCTGTAGGACCAGTAACAAG | This report |
| *pilM* | OMR1 | GGATCCGTTGTTATGGGAATAATTATAGGATTTTTAC | This report |
|  | OMR2 | CTCATTATTTATCACCAACCTAAAGTCCTATTGCCAAAACTTTCCCTCC | This report |
|  | OMR3 | GGAGGGAAAGTTTTGGCAATAGGACTTTAGGTTGGTGATAAATAATGAG | This report |
|  | OMR4 | GTCGACGGTTCTGTAGAATTAACCATTTTCAC | This report |
| *pilN* | OMR9 | GGATCCGAGGAAAAGGTTATTGATGGGAATG | This report |
|  | OMR10 | CTCTTTTATTAATTTTCATTATTTCCTACCCTCTCATTATTTATCACCAACCTAAAG | This report |
|  | OMR11 | CTTTAGGTTGGTGATAAATAATGAGAGGGTAG GAAATAATGAAAATTAATAAAAGAG | This report |
|  | OMR12 | GTCGACCTAATTCATTTTCTATTGTTCCTGTTCC | This report |
| *pilO* | OMR17 | GGATCCCTAAACATGAAAGTAAAGATAGTTCAGGG | This report |
|  | OMR18 | CACACACCTTTTCTATTAGTTTTTAATCCTCTTTTATTAATTTTCATTATTTCCTACCTCC | This report |
|  | OMR19 | GGAGGTAGGAAATAATGAAAATTAATAAAAGAGGATTAAAAACTAATAGAAAAGGTGTGTG | This report |
|  | OMR20 | GTCGACCCTTACACCACTCCCCTTG | This report |
| *pilT* | OAH163 | GTCGACCGACATGTTCATGTTACAATACTTATTAGAATTAG | This report |
|  | OAH146 | GGAGGAGATTTATGCAAAGTTTAGCACGATTATTGATGTA  TTAAATATAGAGGTAATTTG | This report |
|  | OAH147 | GGAGGAGATTTATGCAAAGTTTAGCACGATTATTGATGTA  TTAAATATAGAGGTAATTTG | This report |
|  | OAH148 | GAGCTCCATGAATTAAATAGGTATCCAATATGTGTTCCTC | This report |
| *cpe2280* | OMR25 | GGATCCGACAGTTTCATTAAGTGTAAATGGTAC | This report |
|  | OMR26 | CTTCATGTATAACATATTATCACTCTCCCCCATAGCTTCACACACCTTTTC | This report |
|  | OMR27 | GAAAAGGTGTGTGAAGCTATGGGGGAGAGTGATAATATGTTATACATGAAG | This report |
|  | OMR28 | GTCGACCCACCAGAAATAACTATTCCACTTGC | This report |
| *cpe2279* | OMR33 | GGATCCGCTCAAAGTACTGTTGAACAAATG | This report |
|  | OMR34 | CACCATCTTTCTCTTAATAGTTAGTTTACGTATAACATATTATCACTCTCCTATAAATTCTG | This report |
|  | OMR35 | CAGAATTTATAGGAGAGTGATAATATGTTATACGTAAACTAACTATTAAGAGAAAGATGGTG | This report |
|  | OMR36 | GTCGACCTAGGATCTAAATCTTGTGGCTG | This report |
| *cpe2277* | OMR41 | GGATCCCTGGGAATTTAGATGTTTTAAATAAGG | This report |
|  | OMR42 | GTGATCTTATTTTTCTTTTAGATTAAAATACATCCTTAATCAACTATTTCACCTCCAAC | This report |
|  | OMR43 | GTTGGAGGTGAAATAGTTGATTAAGGATGTATTTTAATCTAAAAGAAAAATAAGATCAC | This report |
|  | OMR44 | GTCGACCTATTCCACCTTCTCTAGCCATAG | This report |
| *cpe1841* | OSRM5 | GTAATACCCTCTGAAAAAATCTTAGAAAAGATAGAAGTATACAGAATG | This report |
|  | OSRM6 | CCTAAATAAATACATAGTTCAATTAATAAATATGCATCTTCCATATCTATATTTCCTCATAAATCACTTAAC | This report |
|  | OSRM7 | GTTAAGTGATTTATGAGGAAATATAGATATGGAAGATGCATATTTATTAATTGAACTATGTATTTATTTAGG | This report |
|  | OSRM8 | CATTTATAACCTTTATTTCATAGTAAGTATCCCTAGAAAGTCCAC | This report |

**Table S2.** Primers for addition of antibody binding tags to BsaA and BsaB

| Purpose | Primer | Sequence (5’-3’) | Reference |
| --- | --- | --- | --- |
| Adding FLAG tag to BsaA Forward | OGC20 | 5’GCAGGTACTAATGCACATAAAGATTACAAGGATGACGATGACAAGTAAGTAATA3’ | This report |
| Adding FLAG tag to BsaA Reverse | OGC21 | 5’TATTACTTACTTGTCATCGTCATCCTTGTAATCTTTATGTGCATTAGTACCTGC3’ | This report |
| Adding HA tag to BsaB Forward | OGC23 | 5’ACCTAACTAAGCGTAGTCTGGGACGTCGTATGGGTATTTAGATAAAATTTCTATATACTTATC3’ | This report |
| Adding HA tag to BsaB reverse | OGC22 | 5’GATAAGTATATAGAAATTTTATCTAAATACCCATACGACGTCCCAGACTACGCTTAGTTAGGT3’ | This report |

**Table S3.** Primers to clone the *bsa* operon into pKRAH1.

| Name | Sequence (5’-3’) | Reference |
| --- | --- | --- |
| OHL145 | CTGCAGGAAAAGGAAGAAAAGGAATAATTATGAAAAAAG | This report |
| OHL146 | GGATCCGTTTTCCATCTTAATTTAATTTTAATATACC | This report |

**Table S4.** Primers for purifying BsaC from *E. coli.*

| Name | Sequence (5’-3’) | Reference |
| --- | --- | --- |
| OGC4 | TTTACACTGCAGTAAAGTTGAAAAGGAAGAAAAGGAATAATTATG | This report |
| OGC5 | TTTAAACCTAGGAATGAAATTTAATTTTTTCTATAATAGTTTTTTATTATAGAA | This report |

**Table S5.** Primers to fuse the FAST gene to the *bsaA* gene.

| Name | Sequence (5’-3’) | Reference |
| --- | --- | --- |
| OSM411 | GTCGACGAAAAGGAAGAAAAGGAATAATTATGAAAAAAGG | This report |
| OSM400 | TGATCCTCCTCCTCCTGATCCTCCTCCTCCTGATTTATGTGCATTAGTACCTGCACCTAAC | This report |
| OSM401 | TCAGGAGGAGGAGGATCAGGAGGAGGAGGATCAATGGAGCATGTTGCCTTTGGCAGTG | This report |
| OSM408 | GGAGATTCATATTGGGTATTTGTAAAAAGAGTATAAGTAATATTTTAAATTAAGTTATAACCATGATGCAAGGAGTGAAAAGG | This report |

**Table S6.** Primers to fuse the FAST gene to the *bsaA* gene.

| Name | Purpose | Sequence (5’-3’) | Reference |
| --- | --- | --- | --- |
| OGC14 | BsaC upstream primer with NdeI sequence for pET24-a expression | CATATGATGAAGAATATAAGAAAAATTTTTTGTG | This report |
| OGC15 | BsaC downstream primer with XhoI sequence for pET24-a expression | CTCGAGATTTAATTTTAATATACCAAAATC | This report |

**Table S7.** Primers to clone the *pilA3* gene for complementation of the in-frame pilA3 mutant.

| Name | Purpose | Sequence (5’-3’) | Reference |
| --- | --- | --- | --- |
| OAH23 | pilA3 N-terminal, SalI | GTCGACCTATTAAGAGAAAGATGGTGATTAAATG | This report |
| OAH127 | pilA3 C-terminal, BamHI | GGATCCAACCCATCTTTTATTTTTTATTTCTAAAAG | This report |

**Supplemental Figure Legends**

**Figure S1.** Immunofluorescence of *C. perfringens* using pre-immune serum (left panel) or serum with anti-PilA3 antibodies. Note the presence of PilA3 at the poles of the cells in the panel on the right.

**Figure S2.** Assay to test for generalized secretion defects in mutants with in-frame deletions of T4P-associated genes with PLC secretion used as the model. PLC secretion causes a precipitation zone to form around the colony on egg yolk agar plates. The indicated strains were patched onto egg yolk agar plates and the zone of precipitation around the colonies was observed. Only the control strain with a *plc* in-frame deletion showed a significant defect in PLC secretion.

**Figure S3.** **(A)** Left panel: Gel-like material produced from expression of the entire *bsa* operon in *C. perfringens* followed by centrifugation of the cells. Right panel: The gel like material in panel A was scraped off and placed in a microfuge tube for visual observation. The amount of material shown is from the growth of 1 L of culture. **(B)** Crystal violet staining of the cells from panel A. Note the diffusely staining material between the bacteria in the left panel. **(C)** Western blot showing TCA precipitation of the supernatants after expression of the *bsaA* operon with a His_6_ tag on the BsaA protein. Note the high molecular weight oligomers present even after heating the sample at 95 °C for 20 min.

**Figure S4.** Western blot showing the dissociation state of oligomers of wild-type BsaA and an N35A mutant after incubation in SDS-PAGE buffer with or without heating at 95 ºC for the times indicated at the top of the figure. Note even though it was incubated for 40 min in SDS-PAGE buffer at room temperature, the N35A mutant was not dissociated in the sample buffer unless heating was applied.

**Figure S5.** Western blot showing boiling cells in 10% SDS for 30 min removes all of the wild-type BsaA oligomers. Cells carrying pSM407 (wild-type), pSM408 (N35A), or pSM409 (K74A) were induced with lactose, pelleted and treated as described. Note that after boiling the BsaA oligomers were no longer present and did not require lysozyme for release, indicating the oligomers were anchored in the membrane of the bacteria.

**Figure S6.** Western blot showing four separate clones of *C. perfringens* carrying pSK2, with an HA tag at the C-terminus of BsaB. Proteins in the culture supernatants were precipitated using TCA and the pellet suspended in SDS-PAGE loading buffer with 100 mM DTT and heated at 95 °C for 20 min. Either 10 or 5 μl of the sample was then loaded onto an SDS-PAGE gel and developed as a western blot.
